# Visualizing BDNF cell-to-cell transfer reveals astrocytes are the primary recipient of neuronal BDNF

**DOI:** 10.1101/255935

**Authors:** Markus A. Stahlberg, Sebastian Kügler, Camin Dean

**Affiliations:** European Neuroscience Institute, Grisebachstrasse 5, 37077 Göttingen, Germany; Center of Nanoscale Microscopy and Physiology of the Brain at Department of Neurology, University Medical Center Göttingen, 37077 Göttingen, Germany

## Abstract

Brain-derived neurotrophic factor (BDNF) is essential for neuronal growth, differentiation, and synaptic plasticity. Although the release and effects of BDNF have been well-studied separately, the transfer of BDNF between cells has not been investigated. Using a four-fluorophore imaging approach to identify both the cell of origin and target cells, we quantified the transfer of BDNF. Surprisingly, we found that astrocytes are the main recipient of neuronally expressed BDNF. We further found that astrocytes specifically take up mature (not pro) BDNF released by neurons. Over-expression of TrkB in neurons redirects released BDNF from astrocytes to neurons, indicating that TrkB levels determine neuronal versus astrocytic BDNF uptake. Increased neuronal activity further increased astrocytic (but not neuronal) uptake of neuronally expressed BDNF. Finally, we demonstrate that astrocytes are not simply a sink for excess BDNF, but that BDNF taken up by astrocytes mediates physiological effects on the astrocytic population by increasing astrocytic territory.

## Introduction

Brain-derived neurotrophic factor (BDNF) was initially discovered and described for its function in mediating neuron survival (Hofer and Barde, 1988; Leibrock et al., 1989), outgrowth of neuronal processes (Lindsay, 1988; McAllister et al., 1995) and neuronal differentiation during development of both the peripheral and central nervous system (Huang and Reichardt, 2001). In addition to its function during development, in the mature central nervous system, BDNF has been identified as a key regulator of synaptic plasticity (Poo, 2001). Its prominent role during development is the reason BDNF knockout mice develop at reduced sizes, live only up to one month and exhibit severe deficits in coordination and balance. They also have recurrent episodes of tonic clonic seizures, and deficits in long-term potentiation (LTP) of synaptic strength underlying learning and memory, most well-studied in the hippocampus (Korte et al., 1998; Patterson et al., 1996). Conversely, exercise increases BDNF, which is implicated in exercise-mediated cognitive improvements (Szuhany et al., 2015).

BDNF is released from dense core vesicles at pre- or post-synaptic (or perisynaptic) sites, constitutively, and in response to neuronal activity (Dean et al., 2009; Dean et al., 2012; Edelmann et al., 2015; Goodman et al., 1996; Hartmann et al., 2001; Kohara et al., 2001; Kojima et al., 2001; Kuczewski et al., 2008; Matsuda et al., 2009). Studies describing the release of BDNF have largely relied on the use of EGFP-tagged BDNF, which (besides creating slightly larger vesicles) is indistinguishable from endogenous BDNF in terms of cellular localization, processing, secretion and binding to its native receptors (Brigadski et al., 2005; Haubensak et al., 1998; Kojima et al., 2001; Matsuda et al., 2009).

Mature BDNF promotes cell survival and the strengthening of synaptic transmission by binding the TrkB receptor (Lu, 2003; Poo, 2001), resulting in increased synaptic vesicle fusion (Gottschalk et al., 1998; Jovanovic et al., 2000; Pozzo-Miller et al., 1999), and post-synaptic AMPA (Caldeira et al., 2007a; Fortin et al., 2012) and NMDA receptor (Caldeira et al., 2007b; Levine et al., 1998) surface expression. Because of its highly basic properties BDNF is thought to act locally following release to affect only synapses in the vicinity of the release site (Poo, 2001). ProBDNF, which is proteolytically cleaved during sorting to produce mature BDNF (Mowla et al., 2001) is also reported to be released (Chen et al., 2005; Yang et al., 2014; Yang et al., 2009), and is implicated in apoptosis and decreasing synaptic transmission by binding the low affinity pan-neurotrophin p75 receptor (Lee et al., 2001; Woo et al., 2005; Yang et al., 2014).

The release of BDNF (assayed by fluorescently-tagged BDNF and ELISA assays), and the effects of BDNF on neurons (using BDNF knockout/knockdown, scavengers, and application of exogenous BDNF) have been well-described. However, transfer of BDNF between cells - in which both cells of origin and target cells can be identified - has not been demonstrated. Transfer of BDNF between connected neurons has long been assumed, beginning with the observation that BDNF protein (but not mRNA) is often visible in anatomical targets of neurons containing BDNF mRNA (Conner et al., 1997; Wetmore et al., 1991; Yan et al., 1997). Transfer is also implied from numerous studies indicating that BDNF is released from neurons (Dean et al., 2009; Dean et al., 2012; Edelmann et al., 2015; Hartmann et al., 2001; Kohara et al., 2001; Kuczewski et al., 2008; Matsuda et al., 2009), and that exogenous radioactively- or fluorescently-labelled BDNF can be taken up by cells (Alderson et al., 2000; Bergami et al., 2008; DiStefano et al., 1992; Rubio, 1997; Santi et al., 2006; Vignoli et al., 2016; von Bartheld et al., 1994). BDNF-GFP fluorescence has also been observed to spread from BDNF-GFP mRNA-injected cells to neighboring neurons (Kohara et al., 2001), and from dendrites of expressing cells to neighboring cell membranes (Kuczewski et al., 2009). But in both studies BDNF-GFP uptake was not examined.

Here we demonstrate BDNF cell-to-cell transfer for the first time. Using a novel live-imaging four-fluorophore approach, we were able to visualize and quantify differential transfer to neurons versus astrocytes. Surprisingly, we found that astrocytes are the major recipient of neuronal BDNF, which was transferred in its mature form. Finally, we demonstrate that the astrocytic population responds physiologically to neuronally released BDNF.

## Results

### BDNF-mRFP1 expressed by neurons is released and transferred to neighboring cells, where it is internalized

When we first expressed a BDNF-mRFP1 construct under the control of the beta actin promotor in dissociated hippocampal cultures transfected, using calcium phosphate, we observed a gradient of RFP fluorescence, highest in the expressing cells, but clearly also present in neighboring cell somata. In following preliminary immunocytochemistry experiments we found that at least some of these neighboring cells were immunopositive for GFAP, an astrocytic cell marker. To exclude low expression of BDNF-mRFP1 in these cells (and identify expressing cells) we designed a construct for targeted expression exclusively in neurons, using a CaMKIIα promotor and joining the BDNF-mRFP1 via a P2A sequence to a second fluorophore (EGFP or ECFP) (**Fig. 1A**). This construct produces two separate peptides: an mRFP1-tagged BDNF and a second cytosolic fluorophore. Transfected cells can then be identified by the cytosolic fluorophore, and the RFP signal in surrounding (non-EGFP expressing) cells corresponds to BDNF-mRFP1 transferred from the transfected cell. By expressing such a construct in dissociated hippocampal cultures, we indeed found that BDNF-mRFP1 was highly abundant in cells that were not transfected. Expression of a control construct, containing cytosolic mRFP1, resulted in RFP fluorescence confined to the expressing cells (**Fig. 1A**).

**Figure 1.**
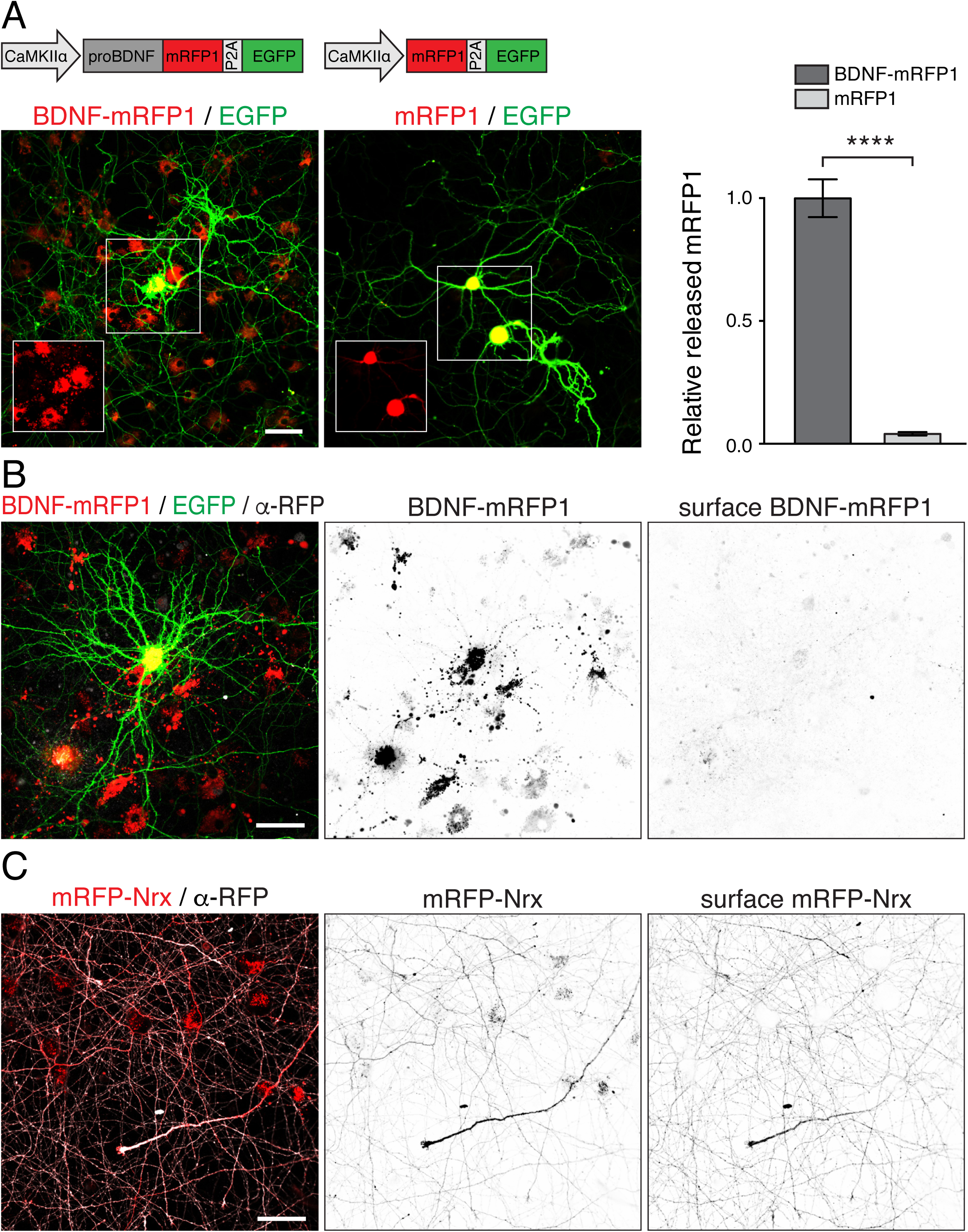
Neuronal expression and transfer of BDNF-mRFP1. (**A**) Construct design for neuronal expression of BDNF-mRFP1-P2A-EGFP (top left) and mRFP1-P2A-EGFP (top middle) with corresponding live cell confocal fluorescence images shown below constructs. Insets show mRFP1 signal of the expressing cell in the indicated region (white box). The mRFP1 fluorophore is not found outside of the EGFP-positive expressing cell, unless it is attached to BDNF; the graph on the right shows normalized released BDNF-mRFP1 and mRFP1 mean fluorescence signal with SEM, from 6 images for each condition (unpaired two- tailed t-test ****p<0.0001 (t=12.36, df=10,)). (**B**) Transferred BDNF-mRFP1 is internalized. Under non-permeabilizing conditions preconjugated primary and secondary antibodies can be used to detect extracellular antigens. Black and white images show the inverted single channel signal of mRFP1 (middle) or the secondary coupled Alexa Fluor 647 fluorophore (right). BDNF-mRFP1 is inaccessible to primary/secondary antibody complexes added to the medium, indicating that BDNF-mRFP1 is internalized. (**C**) Confocal image of a hippocampal culture expressing mRFP-Nrx, where mRFP is present on the extracellular domain. A preconjugated primary/secondary (Alexa Fluor 647) antibody complex detects surface mRFP. Scale bars represent 50 µm.

Because transferred BDNF is not necessarily internalized by recipient cells, but could also reside on the cell surface, we tested for uptake by nearby cells. To do this we immunostained live cultures using an RFP antibody preconjugated to a fluorophore-coupled secondary antibody, under non-permeabilizing conditions, which should detect surface RFP. This antibody complex did not label the characteristic large mRFP1 clusters found in neighboring cell bodies (**Fig. 1B**), suggesting that the BDNF-mRFP1 observed in cells surrounding the transfected neurons is indeed internalized. As a positive control, we used the same antibody complex to immunostain live hippocampal neurons transfected with mRFP-Nrx, where the mRFP is fused to the extracellular domain of neurexin (Nrx). In this case, the RFP antibody signal corresponded to the surface mRFP-Nrx signal (**Fig. 1C**). We can therefore conclude, that BDNF-mRFP1 expressed in neurons, is released and transferred to neighboring cells, where it is internalized.

### BDNF transfer to nearby cells also happens when an EGFP tag or no tag is used

It is curious that a similar transfer of BDNF to neighboring cells has not been reported before, given that fluorophore-tagged BDNF has been used in a great number of studies to examine BDNF localization, trafficking and release. One possible explanation is that transferred BDNF-mRFP1 signal is substantially decreased by application of Ara-C (a pharmacological agent commonly used to limit astrocytic growth), and by fixation and permeabilization, used for performing standard immunocytochemistry experiments (**Suppl. Fig. 1A, B**). Another explanation could be that the majority of previous studies used EGFP tagged BDNF. Because EGFP fluorescence is quenched at low pH, it could be that it is transferred, but the subcellular compartments containing it are acidified, which would mitigate EGFP fluorescence. We tested this hypothesis by generating a construct with interchanged fluorophores (BDNF-EGFP-P2A- mRFP1) and applying 45 mM NH_4_Cl, to de-acidify subcellular compartments. Indeed, under basal conditions only faint EGFP signal could be detected surrounding the expressing cells, but this signal became significantly brighter upon application of NH_4_Cl (**Fig. 2A, B**). The total EGFP mean fluorescence increased by around 54.8%, indicating that BDNF-EGFP is also transferred, but is quenched in subcellular compartments. This signal was specific for transferred BDNF-EGFP; when cytosolic EGFP was expressed under similar conditions, no increase in mean EGFP fluorescence, and no unspecific EGFP signal in cells surrounding EGFP-expressing cells, was detected upon application of NH_4_Cl (**Suppl. Fig. 2A, B**).

**Figure 2.**
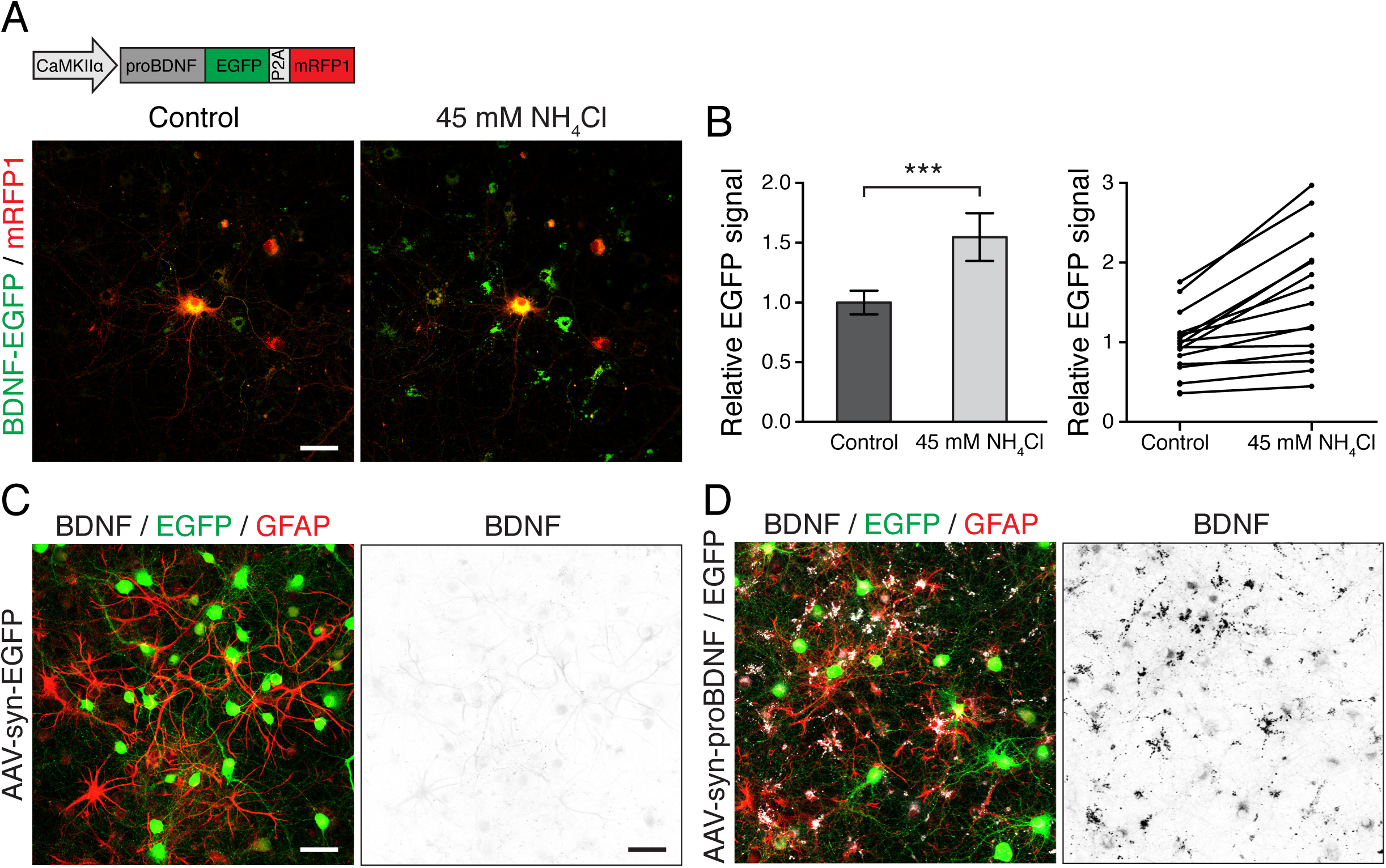
Transfer of BDNF is independent of the mRFP1 fluorophore. (**A**) Detection of transferred BDNF-EGFP is concealed by the quenching of EGFP in acidified subcellular compartments. Exposure of a culture expressing BDNF-EGFP-P2A-mRFP1 (left) to 45 mM NH_4_Cl (right) to de-acidify subcellular compartments, increased EGFP signal allowing the detection of transferred BDNF-EGFP. (**B**) Depiction of the mean EGFP fluorescence with SEM from 5 images each from 3 different cultures (left; paired two-tailed t-test, ***p=0.0003 (t=4.747, df=14)), and the mean EGFP fluorescence of individual images before and after application of NH_4_Cl (right). (**C**) The transfer of BDNF from neurons to astrocytes also occurs with untagged BDNF. Images show BDNF (white) and GFAP (red) immunoreactivity in hippocampal cultures transduced with AAV synapsin-promoter driven EGFP or proBDNF (**D**). Black and white images show the inverted single channel immunoreactivity signal for BDNF. BDNF immunoreactivity is visible under basal conditions and increased when overexpressed in neurons - where transferred BDNF can be found in GFAP immunopositive astrocytes (**D**). The expression pattern and spread of signal is similar for untagged and tagged BDNF, indicating that transfer to neighboring cells is not affected by a fluorophore tag. Scale bars represent 50 µm.

Although we observed transfer of BDNF with two different fluorescent tags, and fluorophores alone were not transferred, this does not exclude the possibility that fluorescent tags might interfere with normal BDNF function. We therefore transduced cultures with AAVs expressing BDNF without a fluorescent tag, and EGFP to mark expressing cells - both driven by the neuron-specific synapsin promoter - and subsequently examined the localization of BDNF using immunocytochemistry (**Fig. 2C, D**). The expression of EGFP was confined to cells with a neuronal morphology and did not overlap with GFAP immunoreactivity, so neuronal expression of BDNF can be inferred. While BDNF immunoreactivity can already be detected under non-overexpressing conditions, it was significantly increased in cultures to which AAV for expression of BDNF was added. Interestingly, the overall pattern of BDNF immunoreactivity was similar to that generated by fluorophore-tagged BDNF: BDNF signal was not only confined to EGFP positive neurons, but also appeared in GFAP-positive cells, implying transfer of untagged BDNF from neurons to astrocytes. Endogenous BDNF was also present in astrocytes in cultures without over-expression of BDNF (**Suppl. Fig. 2C**). Since hippocampal astrocytes lack BDNF mRNA (Ernfors et al., 1990; Rudge et al., 1992), this implies that endogenous BDNF is also transferred from neurons to astrocytes and that BDNF-mRFP1 mimics the behavior of endogenous BDNF in this respect.

### Neuronally-expressed BDNF is predominantly transferred to astrocytes

To determine the relative proportions of BDNF transferred to either neurons or astrocytes, we sparsely transfected neurons with BDNF-mRFP1-P2A-ECFP and utilized AAVs for live-labeling of neighboring neurons and astrocytes. Because immunocytochemistry decreases transferred BDNF signal (**Suppl. Fig. 1A, B**), we used AAV expression of either synapsin promoter-driven LSSmOrange or GFAP2.2 promoter-driven EGFP to quantify differential uptake by neurons or astrocytes, respectively. Neurons can be identified by their LSSmOrange, and astrocytes by their EGFP fluorescence. LSSmOrange (large stokes shift mOrange (Shcherbakova et al., 2012)) is excited by the same wavelength as EGFP, but emits in the orange/red spectrum and is thus separable from ECFP, EGFP and mRFP1 fluorescence by a standard confocal microscope. This allows four-color live cell imaging, where the expression of ECFP identifies transfected neurons, LSSmOrange marks neurons and EGFP marks astrocytes, while BDNF-mRFP1 localization can be quantified, without the need for immunocytochemistry to identify cell types (**Fig. 3A**). We first verified the use of AAVs to label neurons and astrocytes by checking transduction efficiency and specificity of AAV1/2 virions against MAP2 or GFAP immunoreactivity. Around 87.5% of MAP2 positive and 81.9% of GFAP positive cells also expressed the respective fluorophore (**Suppl. Fig. 3**), making this a reliable approach for identifying neurons and astrocytes. We then examined the differential uptake of BDNF-mRFP1 by neurons and astrocytes. Surprisingly, we found that the majority of BDNF-mRFP1 was indeed found in astrocytes rather than neurons (**Fig. 3B**). Only 10.1 ± 7.3% of the released BDNF-mRFP1 was found in neighboring neurons compared to 48.6 ± 6.9% in astrocytes.

**Figure 3.**
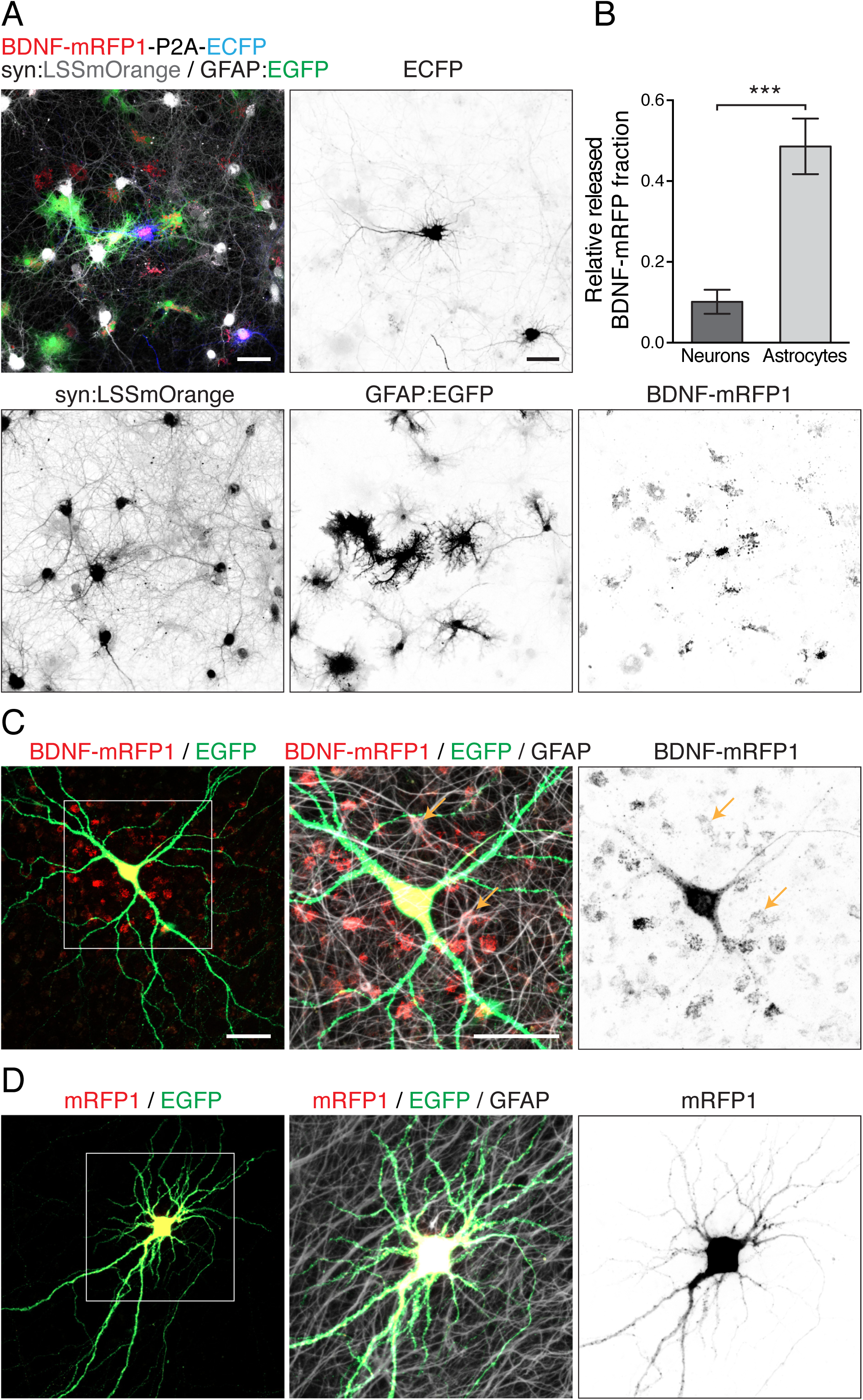
Astrocytes are the major recipient of BDNF released by neurons. (**A**) Live cell image of a hippocampal culture transfected with BDNF-mRFP1-P2A-ECFP and transduced with AAV to express LSSmOrange under synapsin promotor control and EGFP under GFAP2.2 promotor control. Neurons can be identified by LSSmOrange, astrocytes by EGFP, and BDNF-mRFP1 producing cells by ECFP fluorescence. The separate signals are shown in the corresponding black and white images. Much of the BDNF-mRFP1 signal is found in EGFP expressing astrocytes. (**B**) Quantification of 6 images from one culture. Only 10.1 ± 7.3% of the released BDNF-mRFP1 signal was found in neurons, while 48.6 ± 16.8% was found in astrocytes (unpaired two-tailed t-test, ***p=0.0004 (t=5.14, df=10)). The remaining 36.5 ± 10.8% mRFP1 signal did not colocalize exclusively with either area defined by intensity thresholds for LSSmOrange or EGFP. The percents do not sum to 100% because they represent the average of 6 images. (**C**) BDNF neuron-to-astrocyte transfer also occurs in organotypic hippocampal cultures. Images show synapsin promoter driven BDNF-mRFP1- P2A-EGFP or control mRFP1-P2A-EGFP (**D**) gene gun transfected hippocampal organotypic slices, immunostained for GFAP. Signal is shown as average intensity projections of confocal Z-stacks. Left images show EGFP and mRFP1 signal. Images in middle panels show an enlargement of the depicted area in the left image, including the GFAP immunoreactivity signal. The respective single channel image of the mRFP1 signal is shown on the right. Orange arrows indicate transfer and uptake of BDNF-mRFP1 in GFAP immunoreactive cell bodies, which is absent in control mRFP1 expressing cultures. Scale bars represent 50 µm.

While we observed transfer of BDNF from neurons to astrocytes in dissociated hippocampal cultures, this does not necessarily mean that the same occurs under physiological conditions, where the intercellular space is not filled with culture medium or ACSF, but is densely packed with cells and their structures. In order to test this, we prepared organotypic slice cultures of the hippocampus and transfected a few neurons with CaMKIIa- driven BDNF-mRFP1-P2A-EGFP or mRFP1-P2A-EGFP coated gold particles using a gene gun. Subsequently, these slices were fixed and immunostained for GFAP. Indeed, BDNF-mRFP1 signal spread from the neuron of origin and was found in astrocytic cell bodies immunopositive for GFAP (**Fig. 3C**), while cytosolic mRFP1 signal was limited to the neuron of origin (**Fig. 3D**).

### BDNF is transferred to astrocytes in its mature form

Both pro and mature BDNF has been reported to be released. Recently, the participation of astrocytes in the processing and release of previously taken-up proBDNF was proposed(Vignoli et al., 2016). We were therefore curious if the BDNF transferred from neurons to astrocytes that we observed, corresponds to pro or mature BDNF. To answer this question, we made two point mutations in BDNF that render BDNF resistant to intracellular processing, resulting in expression and potential release (Chen et al., 2005; Yang et al., 2014; Yang et al., 2009) of proBDNF only. We incorporated these two point mutations (R129A, R130A) into our BDNF-mRFP1-P2A-ECFP to create cleavage-resistant crBDNF-mRFP1-P2A-ECFP. When we expressed this cleavage-resistant form of BDNF in dissociated hippocampal cultures, we detected no transfer of BDNF-mRFP1 to neighboring cells, compared to the prominent transfer seen for cleavable BDNF (**Fig. 4A**). Thus, the BDNF transferred from neurons to astrocytes corresponds to mature BDNF. This implies that either intracellular processing is required for release of BDNF, or that proBDNF can be released, but astrocytes mainly take up and internalize mature BDNF.

**Figure 4.**
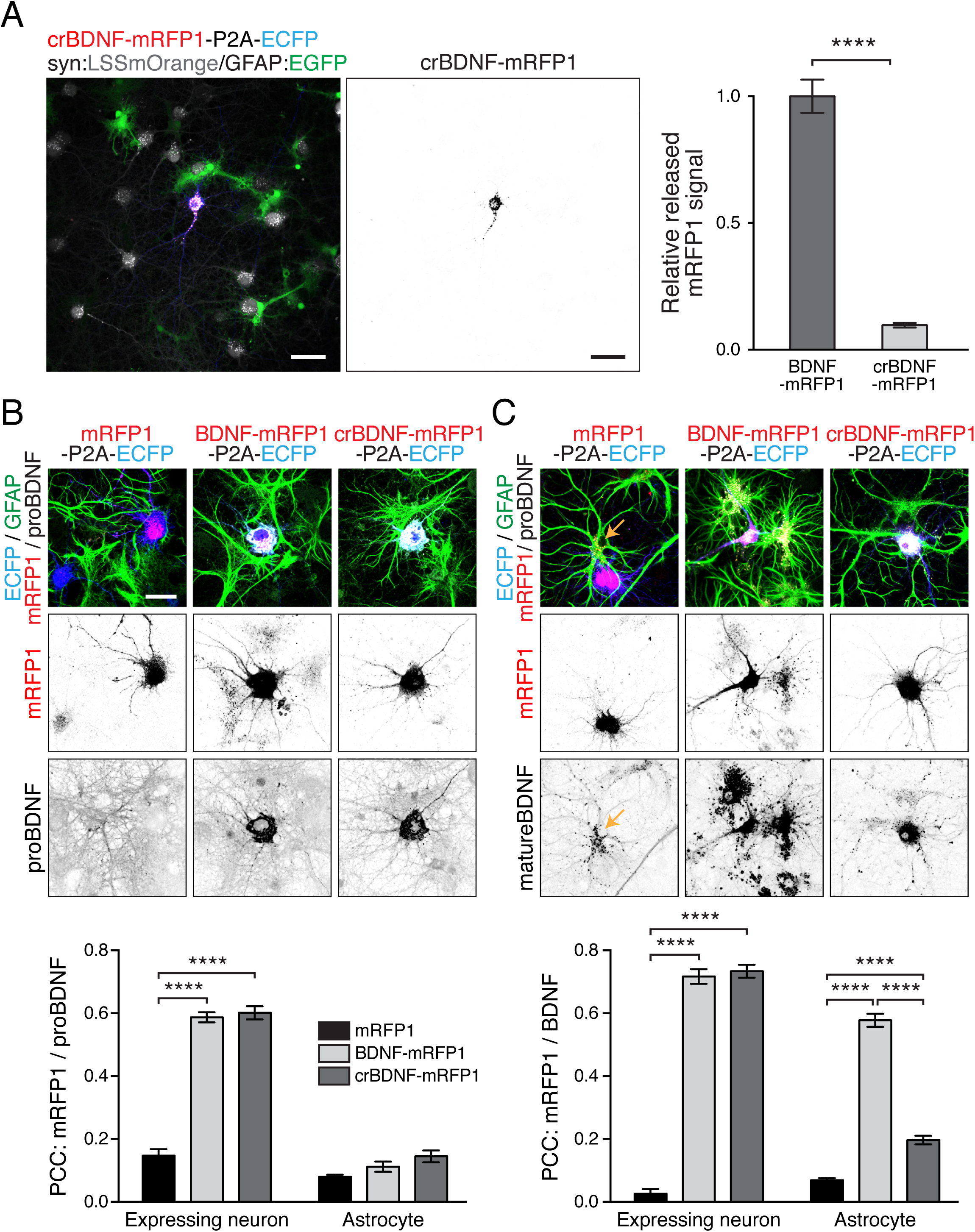
BDNF is transferred in its mature form. (**A**) Confocal image of hippocampal cultures expressing a cleavage-resistant point mutant (R129A, R130A), corresponding to proBDNF, which is not processed to mature BDNF. Surrounding neurons and astrocytes were labeled using AAV promotor-specific expression of LSSmOrange in neurons and EGFP in astrocytes. Inverted single channel image shows crBDNF-mRFP1 fluorescence signal. For each condition, 30 images (5 images per coverslip for 3 cultures) were used to quantify release/transfer of crBDNF-mRFP1. The graph on the right shows BDNF-mRFP1 normalized mean with SEM (unpaired two-tailed t-test, ****p<0.0001 (t=13.64, df=58)). Transfer of cleavage-resistant BDNF is dramatically reduced compared to BDNF. Scale bars represent 50 µm. (**B**) Images of mRFP1, BDNF-mRFP1 and crBDNF-mRFP1 expressing neurons in culture, immunostained for GFAP and for the pro or mature domain (**C**) of BDNF, allowing detection of ProBDNF or all BDNF, respectively. Single channel mRFP1 signal and respective BDNF immunoreactivity are shown in the corresponding black and white images. The two graphs (bottom panels) depict the respective mean Pearson‘s correlation coefficient (PCC) with SEM of mRFP1 signal and BDNF immunoreactivity on expressing neurons or astrocytes from 23 cells per condition from 2 or 3 cultures (two-way ANOVA with Tukey‘s multiple comparisons test, ****p<0.0001 (ProBDNF: F(2, 151)=75.99 and matureBDNF: F(2, 115)=488, both p<0.0001 for the different expression constructs)). ProBDNF immunoreactivity strongly colocalized with BDNF-mRFP1 in expressing neurons, but not astrocytes, while mature BDNF signal colocalized with mRFP1 signal in expressing neurons, but also in astrocytes, indicating that mature, but not proBDNF is transferred to astrocytes. No significant transfer of crBDNF to astrocytes was observed. Note that mature BDNF immunoreactivity in astrocytes is also detected in the absence of BDNF overexpression (**C**, mRFP1-P2A-EGFP, orange arrow). Scale bar represents 30 µm.

To further verify this finding, we expressed either cytosolic mRFP1, BDNF-mRFP1 or crBDNF-mRFP1 (with P2A-ECFP to mark expressing cells) in hippocampal cultures, and immunostained them for GFAP and an antibody that recognizes either the pro or mature domain of BDNF. Increased proBDNF was detected in both BDNF-mRFP1 and crBDNF-mRFP1 expressing neurons, compared to the mRFP1 control, but no increased proBDNF immunoreactivity was found in astrocytes surrounding the expressing cells (**Fig. 4B**). Mature BDNF immunoreactivity was also high in BDNF-mRFP1 and crBDNFmRFP1 expressing neurons, but in contrast was also found (colocalized with BDNF-mRFP1) in surrounding astrocytes (**Fig. 4C**). This substantiates our finding that mature BDNF, and not proBDNF, is transferred from neurons to astrocytes. Notably, under control conditions without overexpression of BDNF, endogenous mature BDNF signal was also high in some astrocytes (**Fig. 4C,** lower left panel), which in the absence of significant proBDNF immunoreactivity of astrocytes, suggests it originates from nearby neurons.

### Transferred BDNF is in late/signaling endosomes and lysosomes (and not in early or recycling endosomes) in both astrocytes and neurons

If we assume that BDNF can be re-released by astrocytes following uptake, as recently proposed (Bergami et al., 2008; Vignoli et al., 2016), we would expect that BDNF that is taken up is sorted to recycling endosomes, from which it can again be released. Alternatively, if astrocytes simply take up excess BDNF to keep extracellular concentrations within a physiological range, BDNF would be expected to be mostly routed to lysosomes. To determine the fate of transferred BDNF, we used electroporation to express EGFP-labeled EEA1, Rab5a, Rab7, Lamp1, LC3, Rab4b or Rab11, in neurons and astrocytes, to mark early and late/signaling endosomes, lysosomes, autophagosomes and recycling endosomes, respectively. In cases where antibodies were available (EEA1, Rab5, Rab7 and Lamp1), the localization of these commonly used marker proteins to their suspected subcellular compartments was verified by immunocytochemistry (**Suppl. Fig. 4**). Following electroporation, cultures were transfected with BDNF-mRFP1-P2A-ECFP and transduced with AAV, to express either synapsin or GFAP2.2 promotor driven LSSmOrange, to mark neurons or astrocytes, respectively. BDNF-mRFP1 colocalization with the respective EGFP-marker protein was then evaluated in ECFP-negative neuronal and astrocytic cell bodies, identified by LSSmOrange fluorescence and morphology (**Fig. 5**). Surprisingly, there was no significant difference in the intracellular localization of internalized BDNF-mRFP1 between neurons and astrocytes. Colocalization, determined by Pearson‘s correlation coefficient, was highest for Rab7 and Lamp1 in both neurons (Rab7=0.55, Lamp1=0.77) and astrocytes (Rab7=0.53, Lamp1=0.75). This suggests a potential similar pathway in neurons and in astrocytes, where the majority of internalized BDNF is localized to late/signaling endosomes and is subsequently degraded, rather than being re-released by recycling endosomes.

**Figure 5.**
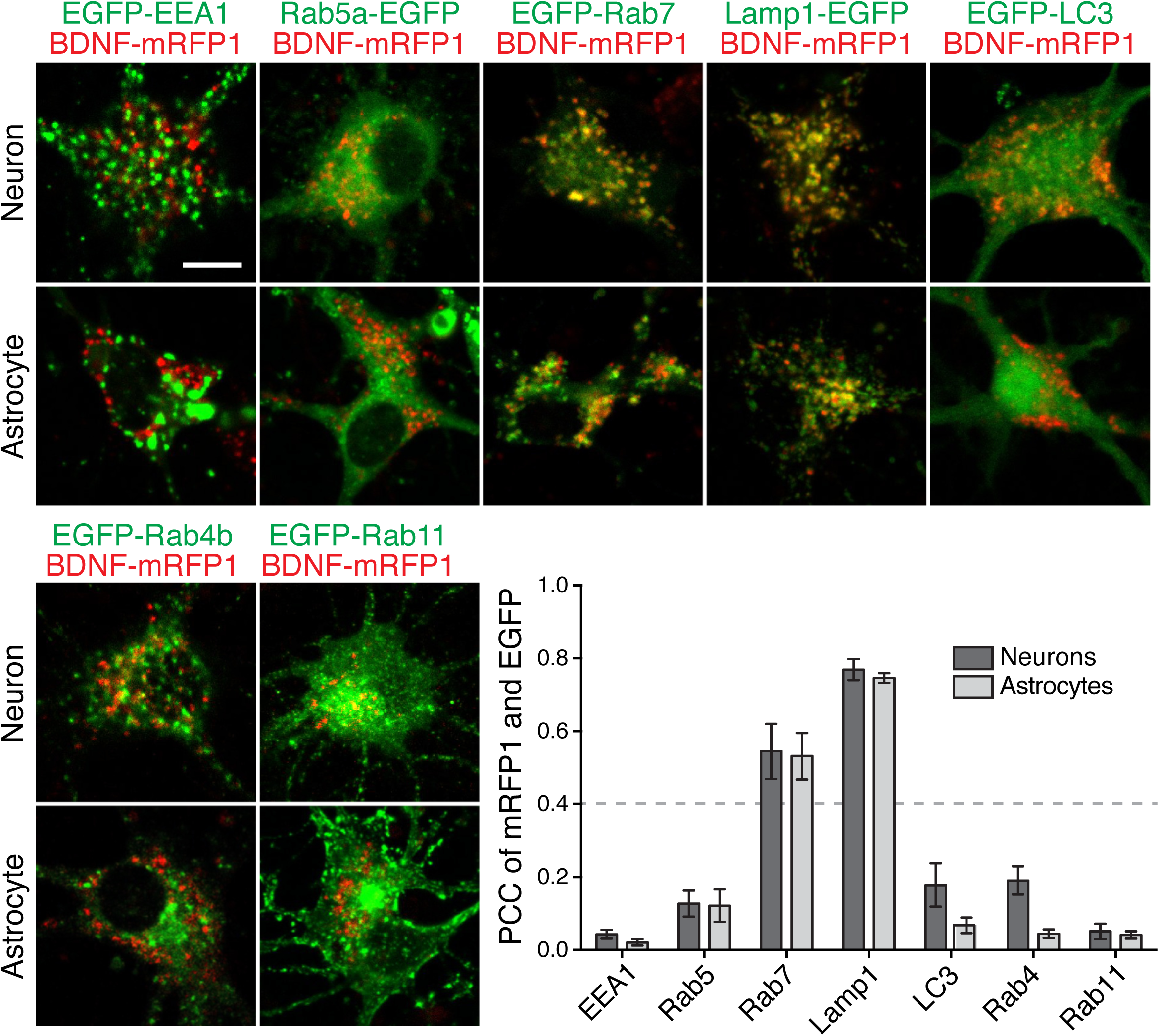
Colocalization of taken-up BDNF-mRFP1 with subcellular organelles. Confocal images of BDNF-mRFP1 and the indicated subcellular marker EGFP fusion proteins in neuronal and astrocytic somata (identified by synapsin or GFAP2.2 driven expression of LSSmOrange and/or morphology). Scale bar represents 10 µm. The graph depicts mean Pearson‘s correlation coefficient (PCC) with SEM, of subcellular markers and BDNF-mRFP1 (n=images of neurons/astrocytes; EEA1 n=15/16, Rab5a n=7/6, Rab7 n=6/6, Lamp1 n=11/8, LC3 n=6/9, Rab4B n=12/24, Rab11 n=10/21). There was no difference in localization of BDNF-mRFP1 taken up by neurons compared to astrocytes. In both cell types taken-up BDNF colocalizes most with Rab7 and Lamp1 in signaling/late endosomes and lysosomes.

### The target receptor for mature BDNF (TrkB) is most abundant on astrocytes

The fact that BDNF is mainly transferred to astrocytes as mature (and not pro) BDNF, prompts the assumption that astrocytes hold more target receptors for mature BDNF than neurons. To test this hypothesis, we examined TrkB immunoreactivity in GFAP-positive astrocytes in BDNF-mRFP1-P2A-ECFP expressing cultures. Astrocytes express the TrkB.T1 receptor (Rose et al., 2003; Rudge et al., 1994), which lacks the intracellular tyrosine kinase domain (Klein et al., 1990), but can be identified by TrkB antibodies, which recognize both full-length and truncated TrkB. We indeed found TrkB signal in GFAP immunopositive cells (**Fig. 6A**). Despite the previously mentioned loss of BDNF-mRFP1 signal through immunocytochemistry, a large proportion of mRFP1 signal overlapped with TrkB signal in astrocytes, although separate mRFP1 and TrkB puncta were also visible. To quantify TrkB signal on neurons and astrocytes, un-transfected cultures were immunostained for MAP2, GFAP and TrkB. Using a threshold for the immunofluorescence signal, combined with manual selection of individual cell bodies, we determined that roughly 38.1% of total TrkB signal was found on astrocytes and only 16.7% was found on neurons (**Fig. 6B**; TrkB signal in areas in which MAP2 and GFAP immunoreactivity overlapped, were not considered). Thus, most TrkB receptors in dissociated hippocampal cultures can be found on astrocytes.

**Figure 6.**
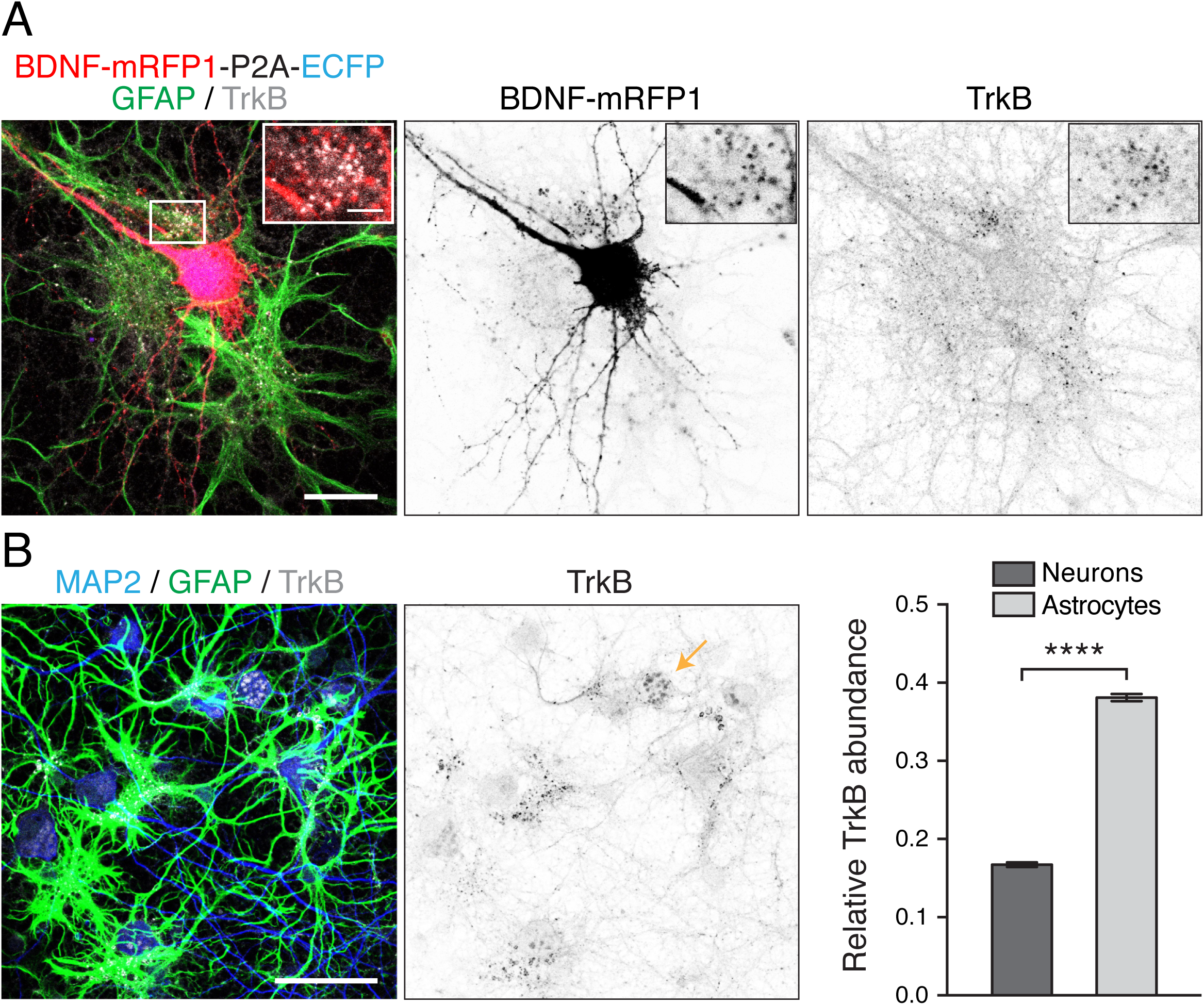
TrkB receptor variants are more abundant on astrocytes than neurons. (**A**) Confocal image of a BDNF-mRFP1-P2A-ECFP overexpressing culture, immunostained for GFAP and TrkB. The indicated area is shown as magnification in the upper right corner. Scale bars represent 25 or 5 µm respectively. Middle and right images show single channel inverted signal of BDNF-mRFP1 or TrkB, including the same magnified area. TrkB appears to be most abundant in astrocytes and TrkB signal prominently co-occurs with BDNF-mRFP1. (**B**) Confocal image of a hippocampal culture immunostained for MAP2, GFAP and TrkB. Middle image depicts TrkB immunoreactivity. The graph on the right shows the relative mean TrkB signal found in either MAP2 or GFAP immunopositive cells with SEM, of 8 images from 12 cultures (n=192, unpaired two-tailed t-test, p****<0.0001 (t=38.48, df=380)). 38.1% of TrkB signal is found in GFAP-positive astrocytes and 16.7% MAP2-positive neurons. The remaining signal could not definitively be assigned to one or the other group, because of MAP2/GFAP signal overlap, signal lower than the selected threshold, or signal in cells positive for neither MAP2 or GFAP. Scale bar represents 50 µm.

### When TrkB is overexpressed in neurons, BDNF is redirected to neurons, and astrocytic territory decreases

Given that most BDNF was found in astrocytes, likely because of their higher TrkB receptor abundance, we next asked if BDNF transfer could be altered by increasing TrkB receptor abundance in neurons. We used AAV mediated expression of synapsin promotor driven cytosolic LSSmOrange or TrkB-LSSmOrange, to label neurons and overexpress the TrkB receptor, and GFAP2.2 promotor driven EGFP to label astrocytes, in BDNF-mRFP1-P2A- ECFP transfected cultures (**Fig. 7A**). To validate the overexpression of TrkB exclusively in neurons, we compared AAV synapsin promotor driven expression of cytosolic EGFP or TrkB-EGFP with MAP2 and GFAP immunoreactivity. AAV mediated expression of syn:EGFP and syn:TrkB-EGFP was indeed confined to MAP2 positive neurons and did not overlap with GFAP positive astrocytes (**Suppl. Fig. 5**). The released BDNF-mRFP1 in both conditions remained the same and was independent of the expression of LSSmOrange or TrkB-LSSmOrange (**Fig. 7B**). However, the amount of BDNF-mRFP1 transferred to neighboring neurons increased from 25.7 to 73.6% of total mRFP1, and the amount of BDNF-mRFP1 transferred to astrocytes decreased from 57.7 to 10.9% (**Fig. 7C**). Thus, increasing TrkB receptor abundance in neurons redirects BDNF-mRFP1 transfer from astrocytes to neurons.

**Figure 7.**
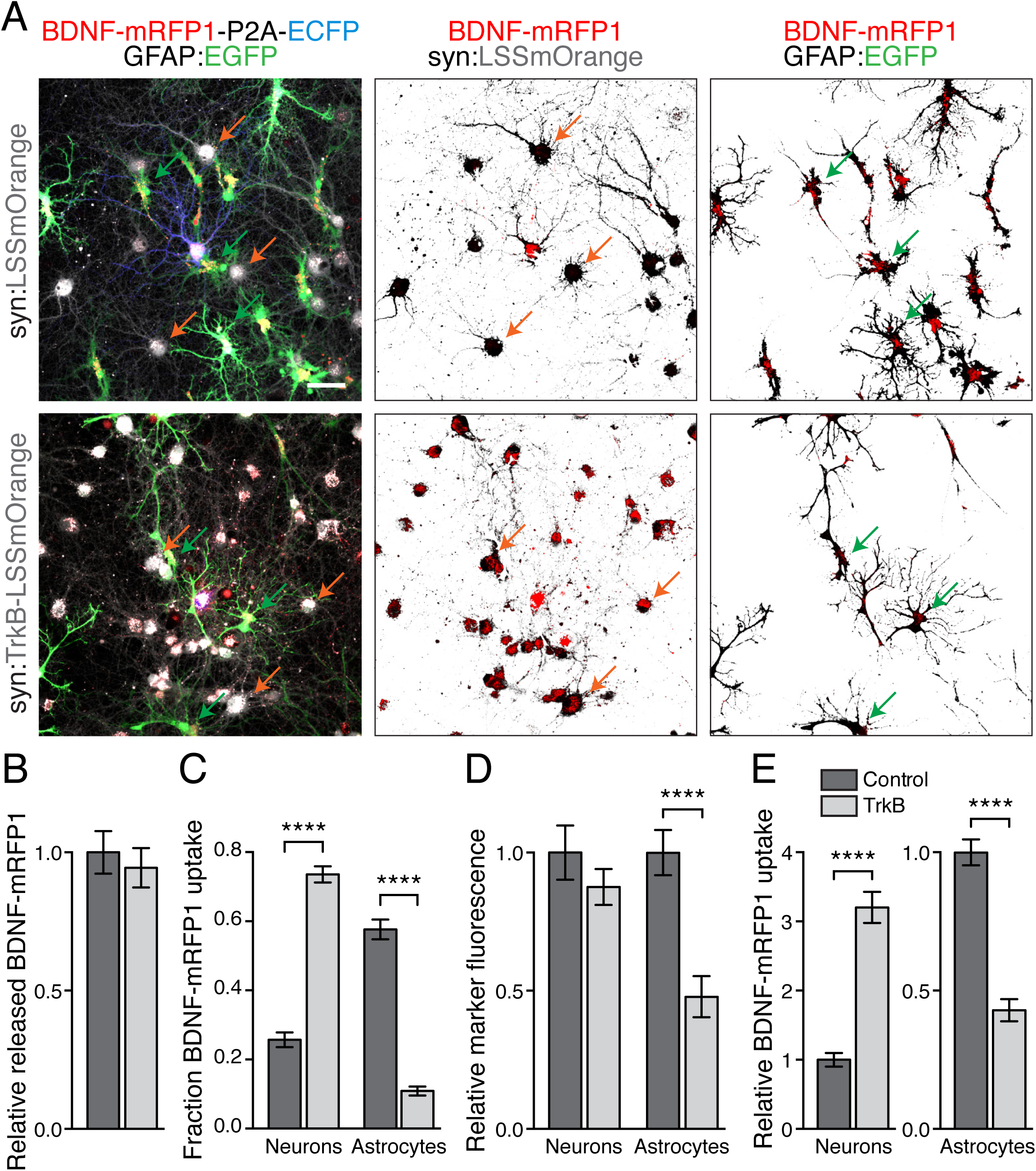
TrkB overexpression in neurons redirects BDNF-mRFP1 to neurons and reduces BDNF-mRFP1 uptake by astrocytes. (**A**) Confocal images of hippocampal cultures expressing BDNF-mRFP1-P2A-ECFP in neurons and transduced with AAV for the expression of GFAP2.2 promoter driven EGFP and synapsin promoter driven LSSmOrange or TrkB-LSSmOrange. BDNF-mRFP1 signal within thresholded and inverted signal of either LSSmOrange-positive neurons (middle) or EGFP-positive astrocytes (right), is also shown for easier visualization of BDNF-mRFP1 found in the respective cell bodies. Selected LSSmOrange and EGFP-positive cells are indicated with yellow or green arrows, respectively. Scale bar represents 50 µm. Under control conditions most mRFP1 signal is found in astrocytes and little is found in neurons. When TrkB is overexpressed in neurons, this is reversed and neurons take up more mRFP1 than astrocytes. (**B**) Quantification of BDNF-mRFP1 released. Graphs represent mean with SEM. There was no difference in the amount of released BDNF-mRFP1 between control and neuronal TrkB overexpression conditions. (**C**) TrkB overexpression in neurons increased the relative amount of BDNF-mRFP1 taken up by neurons from 25.7% to 73.6% and decreased the amount of BDNF- mRFP1 taken up by astrocytes from 57.7% to 10.9%. (**D**) The abundance of astrocytes, measured by surface coverage of EGFP driven by the GFAP2.2 promoter, decreased significantly by 52.2% in cultures in which TrkB was overexpressed in neurons. The increased BDNF-mRFP1 uptake by neurons and decreased uptake by astrocytes persisted when average BDNF-mRFP1 in cells was normalized to cell abundance and total released BDNF-mRFP1 (**E**). Control n=23, TrkB overexpressing n=22; from two cultures. Differences between indicated groups were tested for statistical significance using unpaired two-tailed t-test (**B** (t=0.5257, df=43)**, E** (Neurons: t=9.126, df=43 and Astrocytes: t=9.305, df=43)) and two-way ANOVA (**C** (F(1, 86)=46.69 for cell type and TrkB expression); **D** cell type: F(1, 86)=5.995, p=0.0164 and TrkB expression: F(1, 86)=15.88, p=0.0001)). Statistical differences of individual groups were assessed by Sidak‘s multiple comparisons test (**C, D**),****p<0.0001.

This result was even more striking, when we noticed that in addition to the alteration in relative BDNF-mRFP1 transfer, the relative pixel area covered by EGFP positive astrocytes also decreased by about 52.2%, while the pixel area covered by LSSmOrange expressing cells was unaffected (**Fig. 7D**). Re-directing BDNF away from astrocytes apparently prevents them from extending over a territory that they would otherwise occupy. Because a decreased abundance of astrocytes could potentially account for the decreased BDNF-mRFP1 uptake by astrocytes, we normalized the relative uptake of BDNF-mRFP1 by neurons and astrocytes to released BDNF-mRFP1, and neuronal or astrocytic cell type abundance, determined by LSSmOrange and EGFP signal coverage (**Fig. 7E**). The relative changes are slightly less pronounced after this normalization, but the overall trend was preserved and BDNF-mRFP1 uptake still increased more than threefold in neurons and decreased by 57% in astrocytes in cultures in which TrkB was overexpressed in neurons. Given that decreased BDNF-mRFP1 transferred to astrocytes reduces astrocytic territory, we examined astrocytic territory in experiments in which we expressed cleavage-resistant BDNF, which is not taken up by astrocytes. We found a decrease in astrocytic territory (GFAP2.2 promoter driven EGFP signal) in cultures expressing crBDNF-mRFP1 compared to cultures expressing BDNF- mRFP1 (**Suppl. Fig. 6**). These experiments suggest that an increased abundance of (mature) BDNF released by neurons increases astrocytic territory, and that this effect can be mitigated by decreasing astrocyte accessibility to BDNF.

### Neuronal activity increases BDNF release and uptake by astrocytes, and astrocytic abundance

Because neuronal release of BDNF can occur in an activity-dependent manner(Poo, 2001), we examined the effects of increasing activity using two conditions: 1) a relatively mild stimulation, achieved by increasing the extracellular KCl concentration from 2 mM to 10 mM, and 2) a strong chemical LTP-like stimulation (cLTP), achieved by treatment with bicuculline, glycine, forskolin and rolipram, a protocol derived from inducing LTP in organotypic cultures (Otmakhov et al., 2004; Watt et al., 2004) (**Fig. 8A**). Both stimulations increased the amount of released BDNF-mRFP1; the increase was moderate using KCl stimulation and more pronounced using the cLTP protocol (**Fig. 8B**). Unexpectedly, the increase in release was accompanied by a reduced uptake by neurons, while the amount of BDNF-mRFP1 found in astrocytes increased substantially (**Fig. 8C**).

**Figure 8.**
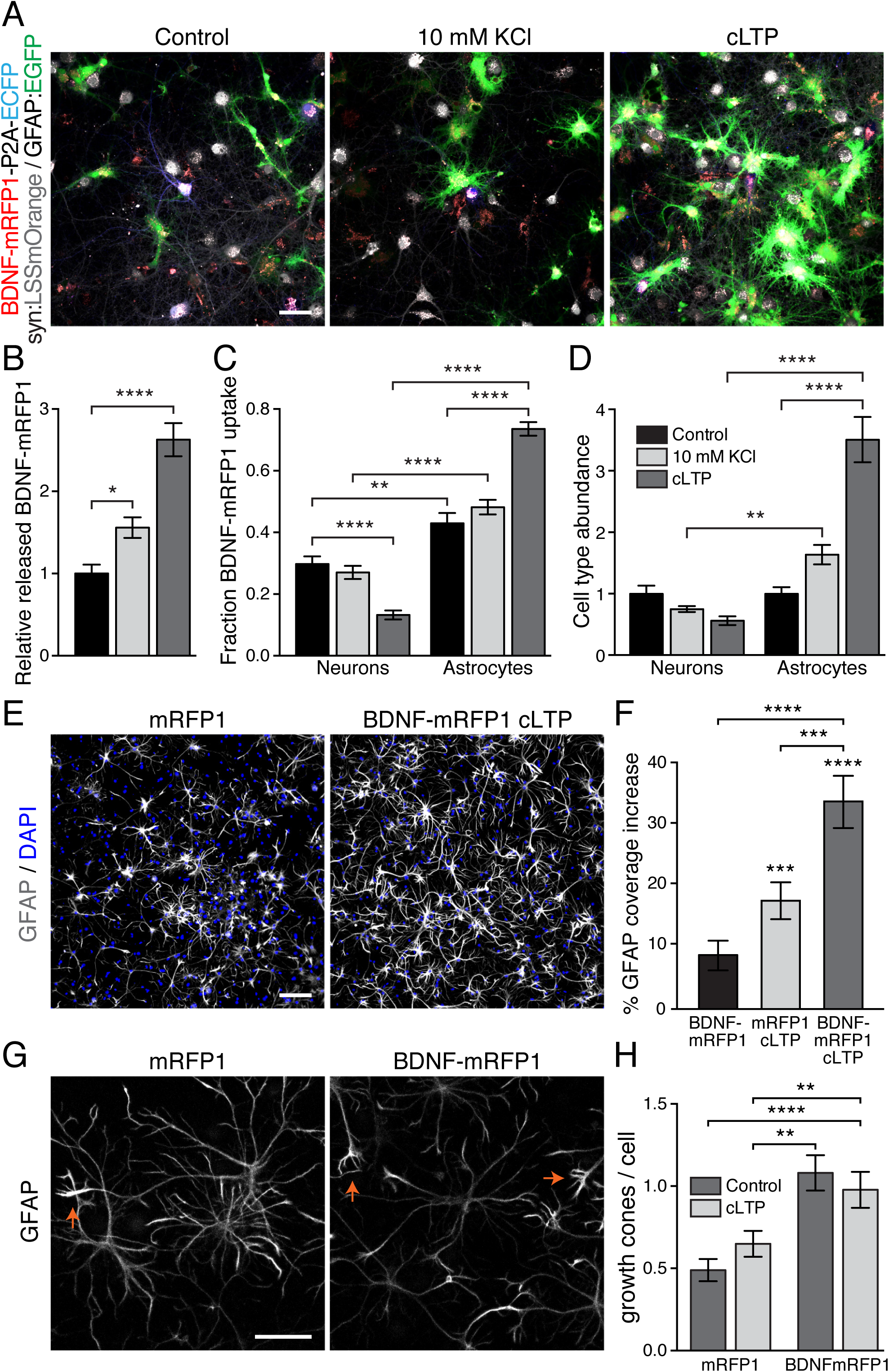
Influence of stimulated BDNF release on astrocytic abundance. (**A**) Confocal images of BDNF-mRFP1-P2A-ECFP expressing neurons in cultures with AAV mediated expression of LSSmOrange in neurons and EGFP in astrocytes, untreated (left) or treated with either 10 mM KCl (middle) or a chemical LTP-like protocol (right). Scale bar represents 50 µm. Graphs of mean fluorescence intensities of released BDNF-mRFP1 with SEM from 5 - 10 images per condition and 3 cultures (control n=25, 10 mM KCl n=30, cLTP n=30). The amount of released BDNF-mRFP1 was increased when cells were stimulated with 10 mM KCl or cLTP. No significant change in relative BDNF-mRFP1 uptake was found for 10 mM KCl stimulation, while cLTP stimulation significantly reduced BDNF-mRFP1 taken up by neurons and increased uptake by astrocytes. (**D**) The increased uptake of BDNF-mRFP1 by astrocytes through cLTP treatment coincided with an increase in astrocytic abundance. Statistical significance was tested using ordinary one-way (**B** F(2, 82)=27.81) or two-way ANOVA (**C** cell type: F(1, 164)=271.3, p<0.0001 and treatment: F(2, 164)=5.201, p=0.0065; **D** cell type: F(1, 164)=68.95 and treatment: F(2, 164)=17.3, both p<0.0001); statistical differences of individual groups was assessed by Dunnett‘s (**B**) or Tukey‘s (**C, D**) multiple comparisons test (*p<0.05, **p<0.01, ****p<0.0001). (**E**) Influence of transferred BDNF abundance on GFAP immunoreactivity. Confocal images show GFAP immunostaining with DAPI fluorescence of cultures expressing mRFP1-P2A-EGFP under control conditions (left) and BDNF-mRFP1-P2A-EGFP stimulated using a chemical LTP-like protocol (right). Scale bar represents 100 µm. Fluorescent GFAP and DAPI pixels above threshold were quantified for 247/6 (mRFP1), 231/6 (mRFP1 + cLTP), 221/5 (BDNF-mRFP1) and 232/5 (BDNF- mRFP1 + cLTP) images/cultures and results shown in the graph on the right (**F**). While there was no difference in the abundance of DAPI-positive pixels (data not shown), GFAP-positive pixel coverage increased with BDNF-mRFP1 expression and cLTP, with the strongest increase when BDNF-mRFP1 expression was combined with cLTP. Results are shown as mean with SEM and normalized to untreated mRFP1-P2A-ECFP. Statistical significance was tested using ordinary two-way ANOVA (BDNF expression: F(1, 928)=16.55 and treatment: F(1, 928)=50.51, both p<0.0001) and statistical differences of individual groups was assessed by Tukey‘s multiple comparisons test (***p<0.001, ****p<0.0001). (**G**) Images of GFAP immunostaining of cultures showing that BDNF-mRFP1 expression in neurons increases the number of astrocytic growth cone like structures compared to mRFP1 expressing neurons in culture. Red arrows indicate exemplary astrocytic growth cone structures. Scale bar represents 50 µm. Astrocytic growth cones were quantified for a total of 90 (mRFP1), 97 (mRFP1/cLTP),87 (BDNF-mRFP1) and 89 (BDNF-mRFP1/cLTP) astrocytes from 8 images each from 3 cultures. Quantification is shown on the right (**H**). Ordinary two-way ANOVA (BDNF expression: F(1, 359)=25.01, p<0.0001 and treatment: F(1, 359)=0.09833, p=0.7540); and statistical differences of individual groups was assessed by Tukey‘s multiple comparisons test (**p<0.01, ****p<0.0001).

We then examined neuronal versus astrocytic area using synapsin-driven LSSmOrange and GFAP2.2 promoter driven EGFP to mark each cell type. While the pixel area covered by neurons decreased insignificantly, the astrocytic territory, identified by EGFP fluorescence pixel area increased approximately 1.6-fold following KCl stimulation and 3.5-fold following cLTP treatment (**Fig. 8D**). To confirm these results independently from AAV-mediated marker expression, we quantified GFAP immunoreactivity in dissociated hippocampal cultures expressing either cytosolic mRFP1 or BDNF-mRFP1, under control conditions and following cLTP treatment (**Fig. 8E, F**) in random regions of entire coverslips (**Suppl. Fig. 7**). We found that GFAP signal pixel area was increased slightly by expression of BDNF-mRFP1 and significantly by cLTP treatment, and increased even more if cLTP was paired with BDNF-mRFP1 expression (**Fig. 8F**). Surprisingly, cLTP treatment alone increased astrocytic territory assayed by GFAP immunoreactivity pixel area, which may be due to the release of endogenous BDNF from neurons and transfer to astrocytes. Note that while GFAP2.2 promoter-driven EGFP labels the entire interior of astrocytes, anti-GFAP labels only fibrous GFAP filaments, thus astrocytic territory may be under-estimated by use of GFAP immunoreactivity. The increase in astrocytic territory is most likely due to an increase in arborization or change in morphology, rather than an increase in astrocyte proliferation, since we did not observe significant changes in number of cells determined by DAPI fluorescence (BDNF-mRFP1 = 1.01 ± 0.16, mRFP1 + cLTP = 1.21 ± 0.17, BDNF-mRFP1 + cLTP = 1.12 ± 0.16; normalized to mRFP1 control) or GFAP levels assayed by western blot (**Suppl. Fig. 8**). In further support of this notion, we found significantly more astrocytic growth cone like structures in cultures in which BDNF levels were elevated by overexpression of BDNF-mRFP1 in neurons (**Fig. 8G, H**).

## Discussion

Our study reveals four important findings. First, we were able to directly observe and quantify the transfer of BDNF from neurons to target cell-types, using a four-color live labeling approach in which both the cell of origin and target cells could be identified. Second, we found that, surprisingly, most BDNF released by neurons is taken up by astrocytes instead of other neurons. Third, using expression of cleavage-resistant proBDNF, we found that only mature BDNF is taken up by astrocytes. And fourth, we discovered that uptake of BDNF by astrocytes affects astrocytic territory.

Given that expression of EGFP-tagged BDNF has been a standard technique for two decades (Haubensak et al., 1998), it is curious that the prominent transfer of BDNF to neighboring cells was not reported before. This is likely due to the quenching of EGFP in acidified subcellular compartments, since de-acidification of subcellular compartments with 45 mM NH_4_Cl revealed robust transfer of BDNF-EGFP to neighboring cells. Importantly, untagged BDNF was also transferred to neighboring cells, excluding possible effects of fluorophore tags. Another possible reason for a lack of previous reports on transfer of BDNF, is the reduction in transferred BDNF-mRFP1 fluorescence following addition of AraC to cultures to limit astroglial proliferation, and following fixation and permeabilization used in immunocytochemistry, which is known to decrease BDNF immunoreactivity (Yan et al., 1997). In addition, the focus of BDNF research is traditionally neurocentric. BDNF transferred to astrocytes was likely observed, but either not identified as such, because a control allowing identification of expressing cells was missing, or simply neglected. BDNF transfer to non-neuronal cells can effectively be seen in recent papers, though it was not noted as such (Andreska et al., 2014; Egan et al., 2003).

We found that astrocytes exclusively take up mature, and not pro, BDNF. This is contrary to a recent report that astrocytes exclusively take up proBDNF secreted by neurons, and release it back to neurons as mature BDNF - based on observations that LTP deficits caused by knockout of the p75 receptor in glial cells could be rescued by expression or addition of mature BDNF (Bergami et al., 2008; Vignoli et al., 2016). In contrast, we found no uptake of proBDNF by astrocytes (despite being able to clearly detect cleavage-resistant proBDNF in expressing neurons) and also found no increase in mature BDNF in neurons following stimulation. We cannot rule out, however, that proBDNF uptake by astrocytes and re-release back to neurons as mature BDNF takes place at levels we cannot detect. In addition, most studies reporting BDNF re-release from astrocytes (Alderson et al., 2000; Bergami et al., 2008; Rubio, 1997; Vignoli et al., 2016) observe it after a large sudden application of recombinant BDNF (Bergami et al., 2008). This could cause unregulated uptake by increased BDNF binding to the low affinity p75 receptor, whereby re-release through recycling endosomes may be increased, simply because more BDNF is present in an unbound or weakly bound state and not retained. It is possible, that in our system we cannot detect such readily re-releasable BDNF, because astrocytes are not exposed to large amounts of exogenous BDNF, and re-release occurs less often than transport of mature BDNF to late/signaling endosomes and lysosomes.

We found that the BDNF-mRFP1 transferred to astrocytes co-localizes with TrkB immunoreactivity, indicating that uptake is likely mediated via the TrkB.T1 receptor. Astrocytic expression of the truncated TrkB.T1 receptor is well documented, while the full-length TrkB receptor is absent and p75 receptor is only expressed in minuscule amounts in astrocytes (Aroeira et al., 2015; Kumar et al., 1993; Roback et al., 1995; Rose et al., 2003; Rudge et al., 1994). Truncated TrkB receptors have been widely thought to primarily limit extracellular BDNF through rapid internalization (Biffo et al., 1995; Fryer et al., 1996), or to act as dominant negatives, controlling full length receptor activation (Eide et al., 1996). But they may signal by non-canonical downstream pathways, through activation of PLC or RhoGDI1 (Aroeira et al., 2015; Ohira et al., 2007; Ohira et al., 2005; Rose et al., 2003). Consistent with this notion, we found that astrocytes do not simply limit extracellular BDNF, but show a physiological response to taken up BDNF by increasing astrocytic territory. This increase is likely caused by changes in astrocytic arborization and surface area rather than proliferation or conversion to GFAP-positive astrocytes, since there was no change in the number of DAPI-stained cells or GFAP protein levels, but there was an increase in astrocytic growth cone-like structures in cultures in which BDNF-mRFP1 was expressed in neurons. This finding is further supported by a previous report that addition of recombinant BDNF to astrocyte monocultures increases cell area via TrkB.T1 mediated Rho GTPase activity (Ohira et al., 2005). In addition, following spinal cord injury, TrkB.T1 knockout astrocytes have slower migration/proliferation in response to BDNF and downregulated migration and proliferation pathways (Matyas et al., 2017). Interestingly, BDNF, TrkB, and astrocytic territory are all increased by exercise (Fahimi et al., 2017), which enhances cognition (Gomez-Pinilla and Hillman, 2013).

Our finding that BDNF is predominantly transferred to astrocytes, where it increases astrocyte territory, prompts a re-evaluation of the previously assumed neuron-neuron effects of BDNF. Astrocytes are well known to affect synapse and circuit function (Volterra and Steinhauser, 2004). Increased astrocytic territory increases synapse number, enhances post- synaptic activity by recruiting AMPA receptors to post-synaptic densities, and enhances presynaptic function by increasing release probability (Chung et al., 2015) - effects that have also been attributed to BDNF, but presumed to result from effects of BDNF directly on neurons. Astrocytic processes are independently regulated and can rapidly extend and retract to interact with synapses in the mouse hippocampus, and these interactions are stabilized at stronger synapses (Haber et al., 2006). BDNF released by neurons and taken up by astrocytes may therefore regulate local astrocyte-synapse interactions to potentiate specific synapses. Given the gradient of BDNF-labelled astrocytes we observe surrounding BDNF-expressing neurons, it is likely that the transfer of BDNF from neurons to astrocytes is not through the medium, but occurs locally through physical contacts that astrocytes make with neurons at tripartite synapses.

In summary, our four-color live labeling approach allowed us to quantify the transfer of neuronal BDNF to neighboring cell-types, and revealed that BDNF is taken up predominantly by astrocytes. It is unlikely that the increased BDNF-mRFP1 we observe in astrocytes compared to neurons, is caused by decreased degradation in astrocytes compared to neurons, given that astrocytes are more efficient at degrading proteins than neurons (Tydlacka et al., 2008). In addition, we found that only mature BDNF is taken up by astrocytes. Finally, we discovered that the uptake of neuronal BDNF by astrocytes does not simply remove excess BDNF, but influences astrocytic territory. It is important to note that although BDNF is not thought to be expressed in mature astrocytes in the hippocampus (Ernfors et al., 1990; Rudge et al., 1992), BDNF is likely in other cell types in the brain in addition to neurons. Cre-dependent removal of BDNF expression in microglia, for example, result in deficits in learning and in learning-dependent synapse formation (Parkhurst et al., 2013). It will be interesting to determine the routes and effects of BDNF transfer between different cell types and brain areas in future studies.

## Methods

### Animals

Use of animals for experimentation was approved and performed according to the specifications of the Institutional Animal Care and Ethics Committees of Göttingen University (T10.31), and of the German animal welfare laws.

### Mammalian expression vectors

Plasmids for the expression of EGFP under synapsin promoter control (pAAV-hsyn-EGFP) and GFAP promoter control (pAAV-GFAP2.2-EGFP), were generated by sub-cloning EGFP under either hSYN or GFAP2.2 promoter control into AAV plasmid vector pTR-UF 22, from which all elements in between the ITRs except the WPRE and the bGH polyadenylation site were removed. The EGFP was replaced by LSSmOrange (pMito-LSSmOrange provided by Vladislav Verkhusha, Addgene #37135) by subcloning to generate pAAV-hsyn-LSSmOrange and pAAV-GFAP2.2-LSSmOrange. A plasmid for the expression of rat proBDNF fused to mRFP1 (pBa-BDNF-mRFP1), originally made by the group of Gary Banker, Ph.D. (Oregon Health & Science University, U.S.A.) was provided by Michael Silverman, Ph.D. (Simon Fraser University, Canada). pEGFP-N1-TrkB (Addgene #32500) was originally produced in the lab of Rosalind Segal, MD, Ph.D. (Dana-Farber Cancer Institute, Harvard Medical School, U.S.A.). TrkB-EGFP was subcloned into the pAAV-hsyn-LSSmOrange backbone, to make pAAV-hsyn- TrkB-EGFP or pAAV-hsyn-TrkB- LSSmOrange. With the assistance of GenScript, proBDNF- mRFP1 was subcloned into pAAV-CaMKIIα-hChR2(T159C)-P2A-EYFP, provided by Karl Deisseroth, MD, Ph.D. (Stanford University, U.S.A.) to generate pAAV-CaMKIIα-BDNF- mRFP1-P2A-EYFP, followed by replacement of EYFP with ECFP (with start codon removed) from pNice-PSD95-CFP, provided by Peter Scheiffele, Ph.D. (University of Basel, Switzerland) to generate pAAV-CaMKIIα-BDNF-mRFP1-P2A-ECFP. Two point mutations (R129A, R130A) were introduced to create cleavage-resistant crBDNF-mRFP1-P2A-ECFP. ECFP was mutated to EGFP, to yield pAAV-CaMKIIα-BDNF-mRFP1-P2A-EGFP. BDNF was removed from the construct to yield pAAV-CaMKIIα-mRFP1-P2A-EGFP and the two fluorophores were swapped to produce pAAV-CaMKIIα-BDNF-EGFP-P2A-mRFP1. CMV promoter-driven expression vectors for GFP-EEA1 and Rab5a-EGFP were provided by Pietro De Camilli, MD (Yale School of Medicine, U.S.A.), GFP-Rab7 was provided by Richard Pagano (Addgene #12605), Lamp1- mGFP was provided by Esteban Dell-Angelica (Addgene #34831), pEGFP-LC3 was provided by Toren Finkel (Addgene #24920). GFP-RAB4B was provided by Gia Voeltz (Addgene #61801) and GFP-rab11 WT was provided by Richard Pagano (Addgene #12674). The pCAG-mRFP-Nrx construct was provided by Peter Scheiffele, Ph.D. (University of Basel, Switzerland). All plasmids were tested by restriction digests and sequenced to verify the correct insert. Sequences and maps were managed with ApE (‘A plasmid Editor’; Wayne Davis, University of Utah, U.S.A.; version 2.0.47).

### Adeno-associated virus (AAV) production

Vectors used to produce AAV serotype 1 & 2 chimeric virions were the pH21 plasmid containing serotype 1, and pRV1 plasmid containing serotype 2 replication and capsid sequences provided by Moritz Rossner, Ph.D. (Psychiatric Clinic of the Ludwig Maximilian University of Munich, Germany) and the adenovirus helper plasmid pFΔ6 was provided by Susanne Schoch McGovern, Ph.D. (University of Bonn Medical Center, Germany). HEK293 cells (Stratagene, cat. no. 240073) at 60% confluence growing in 10 cm dishes were transfected with 10 µg pH21, 10 µg pRV1, 40 µg pFΔ6, and 20 µg expression construct. 48-72 h post-transfection, cells were harvested in 500 µl PBS and lysed by three rounds of freeze-thawing (80 °C for 10 min. followed by 37 °C for 5 min. and vortexing - repeated 3 times). Cell debris was removed by centrifugation at 10,000 rcf for 5 min. The supernatant was collected and stored at 4 °C. 1-2 µl of viral particles were applied per well to dissociated neurons growing in 6-well plates. These chimeric virions successfully transduce both neurons and astrocytes at a transduction efficiency >80. Purified and concentrated AAV serotype 6 virions for the delivery of hsyn-EGFP (2 x 10^8^ vp/µl), hysn-ProBDNF (1.9 x 10^8^ vp/µl), were prepared as described (Toloe et al. Front. Mol. Neurosci., in press), and applied at 10^6^ virions per well to DIV3 neurons growing in 6-well plates.

### Dissociated hippocampal culture

Pregnant Wistar rats were sacrificed between E18 and E19 using CO_2_ followed by cervical dislocation. Embryos were extracted and brains transferred to 10 cm dishes containing Hank‘s Balanced Salt Solution (HBSS) with 10 mM HEPES. Using forceps, hemispheres were separated, meninges removed, and hippocampi extracted and collected in HBSS/HEPES in a 15 ml falcon tube on ice. HBSS was removed and tissue was digested in 2 ml 0.25% Trypsin-EDTA (Gibco) for 20 min at 37 °C, washed 3 times with 5 ml HBSS/HEPES and triturated in 1 ml plating media (DMEM (Dulbecco‘s modified Eagle media) with 10% fetal bovine serum (FBS), 2 mM GlutaMax, and 100 U/ml Penicillin, 100 µg/ml Streptomycin (all from Gibco Thermo Fisher)). Cell suspension was filtered through a 100 µm cell strainer (BD Biosciences) and yield determined by trypan blue exclusion. Cells were plated at 30,500 -37,000 cells/cm^2^ in 600 µl plating medium on 25 mm coverslips (Paul Marienfeld GmbH, cat. no. 0117650) coated with 0.04% polyethylenimine (PEI; Sigma cat. no. P3143) in dH_2_O overnight, followed by 3 washes with dH_2_O, in 6-well plates (CytoOne; cat. no. CC7672-7506), and cultured at 37 °C and 5% CO_2_. The following day plating media was exchanged for 2 ml culture medium (Neurobasal medium with 2% B-27, 2 mM GlutaMax and 100 U/ml Penicillin, 100 µg/ml Streptomycin (all from Gibco)). For the experiments indicated in supplemental figure 1, 5 µM AraC (Sigma) was added once on DIV7. To increase neural network activity, extracellular KCl was increased from 5 mM to 10 mM on DIV3. For chemical LTP-like stimulation, 10 µM bicuculine, 200 µM glycine, 100 nm rolipram and 50 µM forskolin (all from Sigma) were applied once at DIV3-6. Cells were transfected using calcium phosphate transfection and/or transduced with AAV on DIV3. Imaging and immunocytochemistry was performed on DIV14 - 16.

### Calcium phosphate transfection

For each well of a 6-well plate, conditioned medium is collected and replaced with 2 ml pre-equilibrated (37 °C and 5% CO_2_) Opti-MEM (Gibco; cat. no. 11058-021). 60 µl of transfection buffer (274 mM NaCl, 10 mM KCl, 1.4 mM Na_2_HPO_4_, 15 mM glucose, 42 mM HEPES in dH_2_O; pH 7.05 - 7.12, 0.22 µm sterile filtered) is added drop-wise to 60 µl of 7.5 µl 2 M CaCl_2_ and 3.5- 4.0 µg plasmid DNA in dH_2_O) under gentle agitation. This mixture is then incubated for 20 min at room temperature, added drop-wise to Opti-MEM in wells, and cells are returned to the incubator for 90 min. The transfection mix is then replaced with 37 °C and 10% CO_2_ pre-equilibrated Neurobasal medium, incubated for 10 min in the incubator, and Neurobasal medium replaced with conditioned medium.

### Electroporation

Electroporation of subcellular marker proteins (fluorescently-tagged EEA1, Rab5a, Rab7a, Lamp1, LC3, Rab4, and Rab11) was performed using a Rat Neuron Nucleofector Kit (Lonza; cat. no. VPG-1003) and the corresponding Nucleofector 2b Device (Lonza; cat. no. AAB-1001), as recommended by the manufacturer. Dissociated hippocampal cells at a concentration of 3 - 5 x 10^6^ (prior to plating) were pelleted by centrifugation at room temperature and 70 rcf for 5 min, and resuspended in 100 µl Nucleofector solution. 3 µg plasmid DNA was added and the mixture transferred to electroporation cuvettes. For transfection of both neurons and astrocytes, Nucleofector program O-003 (for primary rat hippocampal or cortical neurons and high cell viability) was used. After electroporation, cells were collected in 500 µl pre-equilibrated plating medium (DMEM (Dulbecco‘s modified Eagle media) with 10% fetal bovine serum (FBS), 2 mM GlutaMax, and 100 U/ml Penicillin, 100µg/ml Streptomycin (all from Gibco Thermo Fisher)) and plated at 10^5^ cells/cm^2^ in 600 µl plating medium. Plating medium was exchanged after 6 hours for 2 ml plating medium, which was replaced with culture medium (Neurobasal medium with 2% B-27, 2 mM GlutaMax and 100 U/ml Penicillin, 100 µg/ml Streptomycin (all from Gibco)) the following day.

### Organotypic hippocampal slice culture and Biolistic transfection

P4 Wistar rats were decapitated and brains transferred to 10 cm dishes containing dissection medium (Hank‘s Balanced Salt Solution with 10 mM HEPES, enriched with 6.5 mg/ml D(+)- Glucose). Using forceps and periodontal probes, hemispheres were separated, meninges removed, hippocampi extracted and collected in a separate 10 cm dish containing dissection medium. 400 µm thick coronal slices were prepared using a manual tissue slicer (Stoelting Co. Europe, Dublin, Ireland; cat. no. 51425 equipped with Astra Superior Platinum razor blades). Slices were selected based on their appearance under a stereotactic microscope (Zeiss) and 2 - 3 slices were transferred onto 30 mm hydrophilic polyterafluorethylene membranes with 0.4 µm pores (Millipore; cat. no. PICM0RG50) in 6-well plates filled with 1 ml culture medium (50% Minimal Essential Medium (MEM), 25% Hank‘s Balanced Salt Solution (HBSS), 25% horse serum, 2 mM GlutaMax, 100 U/ml Penicillin, 100 µg/ml Streptomycin (all from Gibco)). Membranes float on top of the medium such that so that slices are provided with nutrients through the medium and oxygen by direct gas exchange with the air. Medium was exchanged the day after dissection and then every 2 - 3 days.

For biolistic transfection of organotypic hippocampal slices, cartridges containing mRFP1-P2A-EGFP and BDNF-mRFP1-P2A-EGFP coated gold particles were made according to the Helios gene gun system (Biorad) instruction manual. 6 - 8 mg of gold was mixed with 100 µl spermidine and sonicated for 20 s. 50 µg of DNA in 50 µl water was added and solution was vortexed briefly. 100 µl of 1 M CaCl_2_ was added drop-wise under gentle vortex, solution sonicated briefly, and incubated for 10 min. at room temp. 7.5 µl of a 20 mg/ml Polyvinylpyrrolidone (PVP) solution was added to 3.5 ml of 100% EtOH. 200 µl of this solution was used to resuspend beads and transfer them to a new tube, and this was repeated to collect all beads in a total volume of approximately 3.2 µl. This solution was drawn into cartridge tubing by syringe, placed on the tubing station and allowed to settle for 2 s, the tubing rotated 90° and allowed to settle for another 2 s, rotated for 5 s, and then rotated while drying under constant CO_2_ flow for 5 min. Tubing was cut into cartridges and bullets ejected through a 50 µm mesh cell strainer (BD Biosciences) using a Helium gas tank with a pressure of 180 PSI, at a distance of 2.5 cm from the slice one day after slice preparation. Slices were then placed back in the incubator and imaged live or following immunocytochemistry 15 - 16 days later.

### Immunocytochemistry, and fixed and live imaging

Cells were fixed with 4 % PFA for 30 min, washed 3 times with PBS (Gibco) and blocked and permeabilized in blocking buffer (2% donkey serum; Sigma, 0.1% Triton X-100; Roth, 0.05% NaN_3_; Roth in 2XPBS) for 30 min. Primary antibody was applied in blocking buffer and incubated overnight at 4 °C. Cells were washed 3 times with PBS, fluorescently labeled secondary antibodies were applied in blocking buffer for 2 h at room temperature, cells were washed 3 more times with PBS, and coverslips were mounted on slides with Fluoromount (Sigma). Antibodies used were: mouse ALDH1L1 (NeuroMab cat. no. 75164), mouse BDNF (Developmental Studies Hybridoma Bank, cat. no. 9), rabbit EEA1, mouse Rab5 and rabbit Rab7 (provided by Reinhard Jahn, Max Planck Institute for Biophysical Chemistry, Göttingen, Germany), guinea pig GFAP (Synaptic Systems, cat. no. 173004), mouse Lamp1 (Enzo Life Sciences cat. no. ADI-VAM-EN001-D), mouse MAP2 (Millipore, cat. no. MAB3418), rat RFP (ChromoTek, cat. no. 5F8), chicken proBDNF (Millipore, cat. no. AB9042), rabbit TrkB (Millipore, cat. no. 07-225). Alexa Fluor 488, 546 and 647 secondary antibodies (Invitrogen) were used.

For live labelling of surface BDNF-mRFP1 and RFP-Nrx, primary RFP antibody was mixed with the respective fluorescently-labeled secondary antibody, at a ratio corresponding to that used for normal immunostaining in a volume of 1 ml, incubated for 30 min at room temperature and applied in fresh culture medium at standard working dilution to the cells which were incubated for another 30 mins before they were transferred to a low profile RC-41LP imaging chamber from Warner Instruments, LLC (Hamden, U.S.A.; cat. no. 64-0368), filled with 1 ml of artificial cerebral spinal fluid (ACSF, in mM: 145.4 NaCl, 5 KCl, 2 CaCl_2_, 1 MgCl_2_, 10 HEPES, 7 D(+)-Glucose) for imaging on a Zeiss A1 TIRF or LSM 710 confocal microscope. For experiments testing acidic quenching of BDNF-EGFP a 90 mM ammonium chloride solution (in mM: 90 NaCl, 5 KCl, 2 CaCl_2_, 2 MgCl_2_, 90 NH_4_Cl, 20 HEPES, 5.5 D(+)-Glucose) was mixed in equal volume with the imaging buffer to reach a final NH_4_Cl concentration of 45 mM.

Fixed and live imaging was conducted using a Zeiss LSM710 confocal microscope, with 405, 458, 488, 561 and 633 nm lasers. For imaging of organotypic hippocampal slices, membranes containing slices were transferred to an imaging slide and kept in imaging buffer between the slide and a cover glass. Slices and live neuronal cultures were imaged using a Plan-APOCHROMAT 10x or 20x objective (Zeiss; cat. no. 420640-9900 and 420650-9901). Fixed samples were imaged with 40x and 63x Plan-APOCHROMAT oil immersion objectives (Zeiss; cat. no. 420762-9800 and 420782-9900). Image acquisition was controlled with Zen Black software.

### Image and data analysis

Image files were analyzed using ImageJ (Wayne Rasband, National Institutes of Health, U.S.A.; version 1.50e) and Fiji (‘Fiji is just ImageJ’, version 2.0.0-rc-43/1.50e). Colocalization was analyzed using the provided Coloc 2 plugin; backprojected pinhole radii and PSF for the 20x W Plan (1332.42 nm & 2D 4.88 µm) and 40x (710.63 nm & 2D 3.75 µm) Plan objectives were calculated using the PSF calculator (Scientific Volume Imaging B.V., Hilversum, Netherlands; https://svi.nl/forms/pinhole.php; https://svi.nl/NyquistCalculator). Quantification of BDNF transfer was done using custom Macros in ImageJ. Fluorescence signal marking the expressing cell, neighboring neurons, or astrocytes was first thresholded to create binary masks for the individual fluorophores. To ensure the detection of whole cell bodies and reduce unspecific noise, a mean filter was applied first. The threshold for the expressing cell was then chosen to incorporate the cell body and primary processes of expressing cells, while thresholds for other neurons and astrocytes was set to incorporate all visible cell bodies, avoiding signal from other fluorophores. Masks for neuronal and astrocytic cells were additionally subtracted from one another to exclude areas of co-occurrence. Total transferred BDNF and uptake by individual cell types was then determined by calculating the average mRFP1 fluorescence found in the respective cell-type area (total mRFP1 signal area minus transfected cell marker area overlapping LSSmOrange or EGFP-positive neurons or astrocytes, respectively). For analysis of astrocytic coverage, GFAP immunostain signal or EGFP fluorescence signal driven by the GFAP2.2 promoter, was thresholded to include all cell bodies and processes. A binary image was created and the number of pixels per image was calculated.

### Statistical analysis

Data calculations were generated using Microsoft Excel software (Microsoft; version 14.6.0) and statistical analysis and representative graphs were created using Prism software (GraphPad; version 6). All reported values in statistical analysis represent the mean, error bars indicate SEM, and Student‘s t-tests are two-tailed type 2, unless otherwise indicated. For all statistical tests, data met the assumptions of the test. All n numbers listed in Figure Legends refer to biological replicates. For t-tests, t-values (t) and degrees of freedom (df) are provided in the corresponding figure legend. For ANOVAs, type, F values, degrees of freedom for numerator (DFn) and denominator (DFd) as F(DFn, DFd) and p values are provided in the corresponding figure legend. To compare individual differences between groups, appropriate corrections for multiple comparisons were applied following GraphPad recommendations. For one-way ANOVAs, Dunnett‘s test for multiple comparisons was used, and for two-way ANOVAs, Sidak‘s or Tukey‘s test for multiple comparisons between two or three conditions, respectively, were used. For all statistical tests, significance is indicated by * p<0.05, ** p<0.01,*** p<0.001, **** p<0.0001.

### Western blotting

Neuronal cultures growing in 6-well plates were harvested in lysis buffer (50 mM Tris-HCl pH 7.5, 150 mM NaCl, 2 mM EDTA, 0.5% NP40, Complete protease inhibitor (Roche)), and analyzed by SDS-PAGE and western blotting to determine GFAP protein levels using guinea pig GFAP (Synaptic Systems, cat. no. 173004) and chicken βIII-tubulin (Abcam, cat. no. ab117716).

## Acknowledgements

We are grateful to Michael Silverman for providing the original BDNF-mRFP1 construct, to Sandra Gebauer for providing molecular biological protocols and HEK cells used in this study, and to Katja Burk for providing assistance with western blotting. This study was supported by a Sofja Kovalevskaja grant from the Alexander von Humboldt Foundation, European Research Council starting grant SytActivity FP7 260916, Deutsche Forschungsgemeinschaft grants CRC889, DE1951, and the Center for Nanoscale Microscopy and Molecular Physiology of the Brain (CNMPB), and the Boehringer Ingelheim Foundation, to C.D.

## Contributions

M.A.S and C.D. designed the study. S.K. provided advice, plasmids and viral vectors. M.A.S. designed and conducted the experiments. M.A.S and C.D. analyzed data, prepared figures and wrote the manuscript. All authors revised and approved the article in its final form.

## Supplemental Figures

**Supplemental Figure 1. Loss of transferred BDNF-mRFP1 fluorescence in culture by standard fixation and permeabilization during immunocytochemistry.** (**A**) Representative images of BDNF-mRFP1-P2A-EGFP or EGFP expressing neurons in different culture conditions or stages of immunocytochemistry (4% PFA fixation and permeabilization). Scale bar represents 50 µm. Inverted images of BDNF-mRFP1 signal are shown below respective fluorescence images. (**B**) Graph represents average released BDNF-mRFP1 fluorescence signal and SEM from 3 - 4 images from 2 - 5 cultures (control/live neuron n=18, AraC n=13, 4% PFA n=18, 4% PFA + permeabilization n=18, 3x PBS wash n=10 and EGFP only n=6). Statistical significance was determined by one-way ANOVA (F(5, 77)=7.006, p<0.0001) and individual differences were determined using Dunnett‘s multiple comparisons test (*p≤0.05,**p≤0.01, ***p≤0.001 and ****p≤0.0001).

**Supplemental Figure 2. NH_4_Cl does not increase EGFP fluorescence in control cytosolic EGFP-expressing cultures.** (**A**) Images show a dissociated hippocampal culture expressing cytosolic EGFP under the control of the synapsin promotor, before (left) and after (right) the application of 45 mM NH_4_Cl. Neither an increase in mean EGFP signal, nor signal in non-expressing cells was detected upon application of NH_4_Cl. (**B**) Graphs show the quantified average EGFP mean signal with SEM (left) and the individual image EGFP mean values (right), before and after the application of NH_4_Cl. (**C**) Image of hippocampal cultures immunostained with GFAP and BDNF. Green arrows indicate GFAP-positive astrocyte cell bodies that also contain BDNF signal. Scale bars represent 50 µm.

**Supplemental Figure 3. AAV-mediated live-labeling of neurons and astrocytes.** Cultures transduced with AAV virions for the expression of EGFP under synapsin promoter control (left) or under GFAP2.2 promotor control (right) and immunostained for MAP2 or ALDH1L1 and GFAP. Quantification is displayed in the graph on the right, showing the mean fraction of transduced cells compared to immunoreactivity of the respective marker, with SEM. EGFP was found in around 87.5 ± 8.2 % (SD) of MAP2 immunopositive neurons and 81.9 ± 11.6 % (SD) of GFAP immunopositive astrocytes (n=6 images for each condition). Only a few cells were positive for ALDH1L1 only and not GFAP. Scale bar represents 25 µm.

**Supplemental Figure 4. Immunocytochemistry validation of subcellular localization of EGFP-tagged marker constructs for EEA1, Rab5a, Rab7 and Lamp1.** Confocal images of hippocampal cultures expressing EGFP-EEA1, Rab5-EGFP, Rab7-EGFP or Lamp1-mGFP immunostained with the corresponding primary antibody. Immunocytochemistry stainings exhibited slightly elevated background signal, but detected the overexpressed subcellular marker proteins. Scale bar represents 25 µm.

**Supplemental Figure 5. Verification of TrkB receptor over-expression exclusively in neurons. (A)** Confocal images of BDNF-mRFP1-P2A-ECFP expressing neurons in cultures transduced with AAV mediating expression of synapsin promoter driven EGFP and immunostained with MAP2, or GFAP (**B**), or transduced with AAV mediating expression of synapsin promoter driven TrkB-EGFP immunostained with MAP2 (**C**), or GFAP (**D**). Synapsin promotor controlled EGFP and TrkB-EGFP can only be detected in MAP2-positive neurons (**A, C**), but not GFAP-positive astrocytes (**B, D**). Scale bar represents 50 µm.

**Supplemental Figure 6. Expression of cleavage resistant BDNF does not increase astrocytic abundance.** Images of BDNF-mRFP1 or crBDNF-mRFP1 expressing neurons in culture, with AAV mediated expression of LSSmOrange in neurons and EGFP in astrocytes. Respective images on the right show inverted single channel images of the GFAP promoter driven EGFP signal. The graph depicts the relative mean EGFP positive pixels per image with SEM. In the cleavage-resistant BDNF-mRFP1 culture astrocyte abundance (EGFP coverage) was significantly lower compared to BDNF-mRFP1 expressing cultures (unpaired two-tailed t-test, ****p≤0.0001 (t=4.366, df=58)). Scale bar represents 50 µm.

**Supplemental Figure 7. GFAP quantification in whole-coverslip images. (A)** Confocal images of tiled whole-coverslip images of hippocampal cultures expressing mRFP1-P2A-EGFP or BDNF-mRFP1-P2A-EGFP in neurons (**B**) treated with a chemical LTP-like stimulation. Magnified images of the area indicated in left images are shown on the right. Cultures show DAPI, GFAP immunoreactivity and EGFP fluorescence signal. Scale bars represent 2500 (left panels) and 250 µm (right panels).

**Supplemental Figure 8. GFAP protein levels in BDNF expressing and stimulated cultures.** Western blot of GFAP cultures in which mRFP1 or BDNF-mRFP1 was expressed in neurons, under control or cLTP treated conditions. Tubulin was used as loading control.

## References

Alderson, R.F., Curtis, R., Alterman, A.L., Lindsay, R.M., and DiStefano, P.S. (2000). Truncated TrkB mediates the endocytosis and release of BDNF and neurotrophin-4/5 by rat astrocytes and schwann cells in vitro. Brain Res 871, 210–222.

Andreska, T., Aufmkolk, S., Sauer, M., and Blum, R. (2014). High abundance of BDNF within glutamatergic presynapses of cultured hippocampal neurons. Front Cell Neurosci 8, 107.

Aroeira, R.I., Sebastiao, A.M., and Valente, C.A. (2015). BDNF, via truncated TrkB receptor, modulates GlyT1 and GlyT2 in astrocytes. Glia 63, 2181–2197.

Bergami, M., Santi, S., Formaggio, E., Cagnoli, C., Verderio, C., Blum, R., Berninger, B., Matteoli, M., and Canossa, M. (2008). Uptake and recycling of pro-BDNF for transmitter-induced secretion by cortical astrocytes. The Journal of cell biology 183, 213–221.

Biffo, S., Offenhauser, N., Carter, B.D., and Barde, Y.A. (1995). Selective binding and internalisation by truncated receptors restrict the availability of BDNF during development. Development 121, 2461–2470.

Brigadski, T., Hartmann, M., and Lessmann, V. (2005). Differential vesicular targeting and time course of synaptic secretion of the mammalian neurotrophins. J Neurosci 25, 7601–7614.

Caldeira, M.V., Melo, C.V., Pereira, D.B., Carvalho, R., Correia, S.S., Backos, D.S., Carvalho, A.L., Esteban, J.A., and Duarte, C.B. (2007a). Brain-derived neurotrophic factor regulates the expression and synaptic delivery of alpha-amino-3-hydroxy-5-methyl-4-isoxazole propionic acid receptor subunits in hippocampal neurons. J Biol Chem 282, 12619–12628.

Caldeira, M.V., Melo, C.V., Pereira, D.B., Carvalho, R.F., Carvalho, A.L., and Duarte, C.B. (2007b). BDNF regulates the expression and traffic of NMDA receptors in cultured hippocampal neurons. Mol Cell Neurosci 35, 208–219.

Chen, Z.Y., Ieraci, A., Teng, H., Dall, H., Meng, C.X., Herrera, D.G., Nykjaer, A., Hempstead, B.L., and Lee, F.S. (2005). Sortilin controls intracellular sorting of brain-derived neurotrophic factor to the regulated secretory pathway. J Neurosci 25, 6156–6166.

Chung, W.S., Allen, N.J., and Eroglu, C. (2015). Astrocytes Control Synapse Formation, Function, and Elimination. Cold Spring Harb Perspect Biol 7, a020370.

Conner, J.M., Lauterborn, J.C., Yan, Q., Gall, C.M., and Varon, S. (1997). Distribution of brain-derived neurotrophic factor (BDNF) protein and mRNA in the normal adult rat CNS: evidence for anterograde axonal transport. J Neurosci 17, 2295–2313.

Dean, C., Liu, H., Dunning, F.M., Chang, P.Y., Jackson, M.B., and Chapman, E.R. (2009). Synaptotagmin-IV modulates synaptic function and long-term potentiation by regulating BDNF release. Nat Neurosci 12, 767–776.

Dean, C., Liu, H., Staudt, T., Stahlberg, M.A., Vingill, S., Buckers, J., Kamin, D., Engelhardt, J., Jackson, M.B., Hell, S.W., and Chapman, E.R. (2012). Distinct Subsets of Syt-IV/BDNF Vesicles Are Sorted to Axons versus Dendrites and Recruited to Synapses by Activity. J Neurosci 32, 5398–5413.

DiStefano, P.S., Friedman, B., Radziejewski, C., Alexander, C., Boland, P., Schick, C.M., Lindsay, R.M., and Wiegand, S.J. (1992). The neurotrophins BDNF, NT-3, and NGF display distinct patterns of retrograde axonal transport in peripheral and central neurons. Neuron 8, 983–993.

Edelmann, E., Cepeda-Prado, E., Franck, M., Lichtenecker, P., Brigadski, T., and Lessmann, V. (2015). Theta Burst Firing Recruits BDNF Release and Signaling in Postsynaptic CA1 Neurons in Spike-Timing-Dependent LTP. Neuron 86, 1041–1054.

Egan, M.F., Kojima, M., Callicott, J.H., Goldberg, T.E., Kolachana, B.S., Bertolino, A., Zaitsev, E., Gold, B., Goldman, D., Dean, M., et al. (2003). The BDNF val66met polymorphism affects activity-dependent secretion of BDNF and human memory and hippocampal function. Cell 112, 257–269.

Eide, F.F., Vining, E.R., Eide, B.L., Zang, K., Wang, X.Y., and Reichardt, L.F. (1996). Naturally occurring truncated trkB receptors have dominant inhibitory effects on brain-derived neurotrophic factor signaling. J Neurosci 16, 3123–3129.

Ernfors, P., Wetmore, C., Olson, L., and Persson, H. (1990). Identification of cells in rat brain and peripheral tissues expressing mRNA for members of the nerve growth factor family. Neuron 5, 511–526.

Fahimi, A., Baktir, M.A., Moghadam, S., Mojabi, F.S., Sumanth, K., McNerney, M.W., Ponnusamy, R., and Salehi, A. (2017). Physical exercise induces structural alterations in the hippocampal astrocytes: exploring the role of BDNF-TrkB signaling. Brain Struct Funct 222, 1797–1808.

Fortin, D.A., Srivastava, T., Dwarakanath, D., Pierre, P., Nygaard, S., Derkach, V.A., and Soderling, T.R. (2012). Brain-derived neurotrophic factor activation of CaM-kinase kinase via transient receptor potential canonical channels induces the translation and synaptic incorporation of GluA1-containing calcium-permeable AMPA receptors. J Neurosci 32, 8127–8137.

Fryer, R.H., Kaplan, D.R., Feinstein, S.C., Radeke, M.J., Grayson, D.R., and Kromer, L.F. (1996). Developmental and mature expression of full-length and truncated TrkB receptors in the rat forebrain. J Comp Neurol 374, 21–40.

Gomez-Pinilla, F., and Hillman, C. (2013). The influence of exercise on cognitive abilities. Compr Physiol 3, 403–428.

Goodman, L.J., Valverde, J., Lim, F., Geschwind, M.D., Federoff, H.J., Geller, A.I., and Hefti, F. (1996). Regulated release and polarized localization of brain-derived neurotrophic factor in hippocampal neurons. Mol Cell Neurosci 7, 222–238.

Gottschalk, W., Pozzo-Miller, L.D., Figurov, A., and Lu, B. (1998). Presynaptic modulation of synaptic transmission and plasticity by brain-derived neurotrophic factor in the developing hippocampus. J Neurosci 18, 6830–6839.

Haber, M., Zhou, L., and Murai, K.K. (2006). Cooperative astrocyte and dendritic spine dynamics at hippocampal excitatory synapses. J Neurosci 26, 8881–8891.

Hartmann, M., Heumann, R., and Lessmann, V. (2001). Synaptic secretion of BDNF after high-frequency stimulation of glutamatergic synapses. EMBO J 20, 5887–5897.

Haubensak, W., Narz, F., Heumann, R., and Lessmann, V. (1998). BDNF-GFP containing secretory granules are localized in the vicinity of synaptic junctions of cultured cortical neurons. Journal of cell science 111 (Pt 11), 1483–1493.

Hofer, M.M., and Barde, Y.A. (1988). Brain-derived neurotrophic factor prevents neuronal death in vivo. Nature 331, 261–262.

Huang, E.J., and Reichardt, L.F. (2001). Neurotrophins: roles in neuronal development and function. Annu Rev Neurosci 24, 677–736.

Jovanovic, J.N., Czernik, A.J., Fienberg, A.A., Greengard, P., and Sihra, T.S. (2000). Synapsins as mediators of BDNF-enhanced neurotransmitter release. Nat Neurosci 3, 323–329.

Klein, R., Conway, D., Parada, L.F., and Barbacid, M. (1990). The trkB tyrosine protein kinase gene codes for a second neurogenic receptor that lacks the catalytic kinase domain. Cell 61, 647–656.

Kohara, K., Kitamura, A., Morishima, M., and Tsumoto, T. (2001). Activity-dependent transfer of brain-derived neurotrophic factor to postsynaptic neurons. Science 291, 2419–2423.

Kojima, M., Takei, N., Numakawa, T., Ishikawa, Y., Suzuki, S., Matsumoto, T., Katoh-Semba, R., Nawa, H., and Hatanaka, H. (2001). Biological characterization and optical imaging of brain-derived neurotrophic factor-green fluorescent protein suggest an activity-dependent local release of brain-derived neurotrophic factor in neurites of cultured hippocampal neurons. Journal of neuroscience research 64, 1–10.

Korte, M., Kang, H., Bonhoeffer, T., and Schuman, E. (1998). A role for BDNF in the late-phase of hippocampal long-term potentiation. Neuropharmacology 37, 553–559.

Kuczewski, N., Porcher, C., Ferrand, N., Fiorentino, H., Pellegrino, C., Kolarow, R., Lessmann, V., Medina, I., and Gaiarsa, J.L. (2008). Backpropagating action potentials trigger dendritic release of BDNF during spontaneous network activity. J Neurosci 28, 7013–7023.

Kuczewski, N., Porcher, C., Lessmann, V., Medina, I., and Gaiarsa, J.L. (2009). Activity-dependent dendritic release of BDNF and biological consequences. Molecular neurobiology 39, 37–49.

Kumar, S., Pena, L.A., and de Vellis, J. (1993). CNS glial cells express neurotrophin receptors whose levels are regulated by NGF. Brain Res Mol Brain Res 17, 163–168.

Lee, R., Kermani, P., Teng, K.K., and Hempstead, B.L. (2001). Regulation of cell survival by secreted proneurotrophins. Science 294, 1945–1948.

Leibrock, J., Lottspeich, F., Hohn, A., Hofer, M., Hengerer, B., Masiakowski, P., Thoenen, H., and Barde, Y.A. (1989). Molecular cloning and expression of brain-derived neurotrophic factor. Nature 341, 149–152.

Levine, E.S., Crozier, R.A., Black, I.B., and Plummer, M.R. (1998). Brain-derived neurotrophic factor modulates hippocampal synaptic transmission by increasing N-methyl-D-aspartic acid receptor activity. Proc Natl Acad Sci U S A 95, 10235–10239.

Lindsay, R.M. (1988). Nerve growth factors (NGF, BDNF) enhance axonal regeneration but are not required for survival of adult sensory neurons. J Neurosci 8, 2394–2405.

Lu, B. (2003). BDNF and activity-dependent synaptic modulation. Learn Mem 10, 86–98.

Matsuda, N., Lu, H., Fukata, Y., Noritake, J., Gao, H., Mukherjee, S., Nemoto, T., Fukata, M., and Poo, M.M. (2009). Differential activity-dependent secretion of brain-derived neurotrophic factor from axon and dendrite. J Neurosci 29, 14185–14198.

Matyas, J.J., O’Driscoll, C.M., Yu, L., Coll-Miro, M., Daugherty, S., Renn, C.L., Faden, A.I., Dorsey, S.G., and Wu, J. (2017). Truncated TrkB.T1-Mediated Astrocyte Dysfunction Contributes to Impaired Motor Function and Neuropathic Pain after Spinal Cord Injury. J Neurosci 37, 3956–3971.

McAllister, A.K., Lo, D.C., and Katz, L.C. (1995). Neurotrophins regulate dendritic growth in developing visual cortex. Neuron 15, 791–803.

Mowla, S.J., Farhadi, H.F., Pareek, S., Atwal, J.K., Morris, S.J., Seidah, N.G., and Murphy, R.A. (2001). Biosynthesis and post-translational processing of the precursor to brain-derived neurotrophic factor. J Biol Chem 276, 12660–12666.

Ohira, K., Funatsu, N., Homma, K.J., Sahara, Y., Hayashi, M., Kaneko, T., and Nakamura, S. (2007). Truncated TrkB-T1 regulates the morphology of neocortical layer I astrocytes in adult rat brain slices. Eur J Neurosci 25, 406–416.

Ohira, K., Kumanogoh, H., Sahara, Y., Homma, K.J., Hirai, H., Nakamura, S., and Hayashi, M. (2005). A truncated tropomyosin-related kinase B receptor, T1, regulates glial cell morphology via Rho GDP dissociation inhibitor 1. J Neurosci 25, 1343–1353.

Otmakhov, N., Tao-Cheng, J.H., Carpenter, S., Asrican, B., Dosemeci, A., Reese, T.S., and Lisman, J. (2004). Persistent accumulation of calcium/calmodulin-dependent protein kinase II in dendritic spines after induction of NMDA receptor-dependent chemical long-term potentiation. J Neurosci 24, 9324–9331.

Parkhurst, C.N., Yang, G., Ninan, I., Savas, J.N., Yates, J.R., 3rd, Lafaille, J.J., Hempstead, B.L., Littman, D.R., and Gan, W.B. (2013). Microglia promote learning-dependent synapse formation through brain-derived neurotrophic factor. Cell 155, 1596–1609.

Patterson, S.L., Abel, T., Deuel, T.A., Martin, K.C., Rose, J.C., and Kandel, E.R. (1996). Recombinant BDNF rescues deficits in basal synaptic transmission and hippocampal LTP in BDNF knockout mice. Neuron 16, 1137–1145.

Poo, M.M. (2001). Neurotrophins as synaptic modulators. Nature reviews Neuroscience 2, 24–32.

Pozzo-Miller, L.D., Gottschalk, W., Zhang, L., McDermott, K., Du, J., Gopalakrishnan, R., Oho, C., Sheng, Z.H., and Lu, B. (1999). Impairments in high-frequency transmission, synaptic vesicle docking, and synaptic protein distribution in the hippocampus of BDNF knockout mice. J Neurosci 19, 4972–4983.

Roback, J.D., Marsh, H.N., Downen, M., Palfrey, H.C., and Wainer, B.H. (1995). BDNF-activated signal transduction in rat cortical glial cells. Eur J Neurosci 7, 849–862.

Rose, C.R., Blum, R., Pichler, B., Lepier, A., Kafitz, K.W., and Konnerth, A. (2003). Truncated TrkB-T1 mediates neurotrophin-evoked calcium signalling in glia cells. Nature 426, 74–78.

Rubio, N. (1997). Mouse astrocytes store and deliver brain-derived neurotrophic factor using the non-catalytic gp95trkB receptor. Eur J Neurosci 9, 1847–1853.

Rudge, J.S., Alderson, R.F., Pasnikowski, E., McClain, J., Ip, N.Y., and Lindsay, R.M. (1992). Expression of Ciliary Neurotrophic Factor and the Neurotrophins-Nerve Growth Factor, Brain-Derived Neurotrophic Factor and Neurotrophin 3-in Cultured Rat Hippocampal Astrocytes. Eur J Neurosci 4, 459–471.

Rudge, J.S., Li, Y., Pasnikowski, E.M., Mattsson, K., Pan, L., Yancopoulos, G.D., Wiegand, S.J., Lindsay, R.M., and Ip, N.Y. (1994). Neurotrophic factor receptors and their signal transduction capabilities in rat astrocytes. Eur J Neurosci 6, 693–705.

Santi, S., Cappello, S., Riccio, M., Bergami, M., Aicardi, G., Schenk, U., Matteoli, M., and Canossa, M. (2006). Hippocampal neurons recycle BDNF for activity-dependent secretion and LTP maintenance. EMBO J 25, 4372–4380.

Shcherbakova, D.M., Hink, M.A., Joosen, L., Gadella, T.W., and Verkhusha, V.V. (2012). An orange fluorescent protein with a large Stokes shift for single-excitation multicolor FCCS and FRET imaging. J Am Chem Soc 134, 7913–7923.

Szuhany, K.L., Bugatti, M., and Otto, M.W. (2015). A meta-analytic review of the effects of exercise on brain-derived neurotrophic factor. J Psychiatr Res 60, 56–64.

Tydlacka, S., Wang, C.E., Wang, X., Li, S., and Li, X.J. (2008). Differential activities of the ubiquitin-proteasome system in neurons versus glia may account for the preferential accumulation of misfolded proteins in neurons. J Neurosci 28, 13285–13295.

Vignoli, B., Battistini, G., Melani, R., Blum, R., Santi, S., Berardi, N., and Canossa, M. (2016). Peri-Synaptic Glia Recycles Brain-Derived Neurotrophic Factor for LTP Stabilization and Memory Retention. Neuron.

Volterra, A., and Steinhauser, C. (2004). Glial modulation of synaptic transmission in the hippocampus. Glia 47, 249–257.

von Bartheld, C.S., Kinoshita, Y., Prevette, D., Yin, Q.W., Oppenheim, R.W., and Bothwell, M. (1994). Positive and negative effects of neurotrophins on the isthmo-optic nucleus in chick embryos. Neuron 12, 639–654.

Watt, A.J., Sjostrom, P.J., Hausser, M., Nelson, S.B., and Turrigiano, G.G. (2004). A proportional but slower NMDA potentiation follows AMPA potentiation in LTP. Nat Neurosci 7, 518–524.

Wetmore, C., Cao, Y.H., Pettersson, R.F., and Olson, L. (1991). Brain-derived neurotrophic factor: subcellular compartmentalization and interneuronal transfer as visualized with anti-peptide antibodies. Proc Natl Acad Sci U S A 88, 9843–9847.

Woo, N.H., Teng, H.K., Siao, C.J., Chiaruttini, C., Pang, P.T., Milner, T.A., Hempstead, B.L., and Lu, B. (2005). Activation of p75NTR by proBDNF facilitates hippocampal long-term depression. Nat Neurosci 8, 1069–1077.

Yan, Q., Rosenfeld, R.D., Matheson, C.R., Hawkins, N., Lopez, O.T., Bennett, L., and Welcher, A.A. (1997). Expression of brain-derived neurotrophic factor protein in the adult rat central nervous system. Neuroscience 78, 431–448.

Yang, J., Harte-Hargrove, L.C., Siao, C.J., Marinic, T., Clarke, R., Ma, Q., Jing, D., Lafrancois, J.J., Bath, K.G., Mark, W., et al.(2014). proBDNF negatively regulates neuronal remodeling, synaptic transmission, and synaptic plasticity in hippocampus. Cell Rep 7, 796–806.

Yang, J., Siao, C.J., Nagappan, G., Marinic, T., Jing, D., McGrath, K., Chen, Z.Y., Mark, W., Tessarollo, L., Lee, F.S., et al. (2009). Neuronal release of proBDNF. Nat Neurosci 12, 113–115.

